# SUGARCANE’S DROUGHT MEMORY LEGACY: HOW PAST STRESS SHAPES FUTURE RESILIENCE

**DOI:** 10.64898/2026.09.29.755288

**Authors:** Maria D. Pissolato, Tamires S. Martins, Marcela T. Miranda, Gabriel S. Pires, Rafael L. Almeida, Larissa P. Cruz, Leonardo P. Sousa, Adilson P. Domingues-Jr, Eduardo C. Machado, Alisdair Fernie, Rafael V. Ribeiro

**Affiliations:** Laboratory of Crop Physiology (LCroP), Department of Plant Biology, Institute of Biology, State University of Campinas (UNICAMP), Campinas SP, Brazil; Center for Plant Molecular Breeding (CeM2P), Institute of Biology, State University of Campinas (UNICAMP), Campinas, SP, Brazil; Research Group on Plant Biology under Mediterranean Conditions, Universitat de les Illes Balears (UIB) – Instituto de Investigaciones Agroambientales y de Economía del Agua (INAGEA), Palma, Illes Balears, Spain; Institute of Agricultural and Environmental Sciences, Estonian University of Life Sciences, Tartu, Estonia; Central Metabolism, Max Planck Institute of Molecular Plant Physiology, Potsdam, Germany

**Keywords:** Developmental stage, water use efficiency, primary metabolism, *saccharum* spp., somatic stress memory

## Abstract

Plants frequently experience recurrent drought events separated by periods of rehydration. Although drought imposes strong constraints on plant physiology, prior exposure may alter subsequent stress responsiveness through memory-based mechanisms. Here, we investigated whether recurrent drought at distinct developmental stages establishes stress memory in sugarcane and whether this response persists across vegetative propagation. Two genotypes contrasting in drought tolerance and productivity (IACCTC07-8008 and IACSP95-5000, respectively) were grown under greenhouse conditions and subjected to three drought cycles imposed either at tillering or maturation stage. Gas exchange, photochemical performance, leaf water status, primary metabolite profile, and growth traits were assessed across cycles, and vegetative propagules were subsequently evaluated under renewed drought. The first drought cycle imposed strong limitations on carbon assimilation and photochemistry in both genotypes and developmental stages. However, subsequent cycles resulted in attenuated reductions in *A* and *g*, improved intrinsic water use efficiency, and partial stabilization of PSII performance, indicating a modified stress-response trajectory. Young plants displayed earlier improvements (from the second cycle), whereas in mature plants this shift was evident mainly during the third cycle. Recurrent drought promoted sustained reorganization of amino acid, carbohydrate, organic acid and polyol metabolism alongside increased root biomass and higher relative water content during later cycles. Importantly, propagules derived from drought-conditioned plants exhibited faster recovery of photosynthetic performance and reduced cumulative physiological impairment under renewed drought, despite showing similar stress sensitivity at maximum water deficit. This persistence of enhanced recovery capacity across vegetative propagation indicates that drought-induced memory was maintained beyond the initially stressed plants. Together, our findings demonstrate that recurrent drought establishes a metabolically imprinted state in clonal sugarcane, integrating physiological adjustment, metabolic reprogramming, and whole-plant acclimation. These results highlight the potential of stress memory as a mechanism supporting resilience in perennial crops exposed to increasingly recurrent drought events.

**Highlights:**

- Repeated drought cycles enhance photosynthetic performance and water use efficiency.
- Drought memory develops faster at tillering than at the maturation stage.
- Metabolomic shifts reveal central carbon and amino acid reorganization under drought.
- Stress-induced metabolic imprint persists in clonally derived propagules.

## 1. Introduction

Increasing climatic variability is intensifying the frequency and severity of drought events, posing a major threat to global agricultural productivity (Koua et al., 2021; Rajak, 2021). This scenario is expected to worsen in the coming decades due to rising temperatures and changes in precipitation patterns (Wang et al., 2014). Sugarcane is widely cultivated in tropical and subtropical regions, particularly in Brazil, China, India, and Mexico (Bordonal et al., 2018), and, as a perennial crop, it is frequently exposed to seasonal water deficits throughout its growth cycle, with significant consequences for plant growth and productivity (Qin et al., 2023; Ribeiro et al., 2013).

Drought stress can occur at any stage of plant development and affects plant water relations across multiple levels, including whole-plant, organ, cellular, and molecular scales (Li et al., 2014; Muscolo et al., 2015). Among the earliest responses to drought, physiological alterations are the initial responses exhibited by plants (Bechtold et al., 2016). Partial stomatal closure occurs in response to water deficit. Since the influx of CO_2_ and transpiration share the same gateway, there is an intrinsic trade-off between reducing water loss through transpiration and the ability to assimilate CO_2_, consequently reducing plant growth and biomass (Ribeiro et al., 2013; Atkinson et al., 2016). Under moderate drought, declines in photosynthetic rate are primarily driven by diffusive limitations, including stomatal and mesophyll conductance, whereas biochemical impairments of the photosynthetic machinery become more pronounced under severe water deficit (Atkinson et al., 2016; Chaves and Oliveira, 2004), that often results in reduced investment in shoot growth and increased allocation to the root system at the expense of aboveground biomass (Bechtold et al., 2016; Fleta-Soriano and Munné-Bosch, 2016). Additional adjustments include the accumulation of compatible solutes, reductions in xylem hydraulic conductivity and leaf water content and increased production of antioxidant compounds across different plant organs (Choat et al., 2012; Marcos et al., 2018a; Pissolato et al., 2020).

Recent studies suggest that the capacity of plants to modulate their response mechanisms under continuous stress is a major determinant of their future fitness and underpins their ability to persist across diverse habitats (Fleta-Soriano and Munné-Bosch, 2016; Liu et al., 2021). Within this context, the concept of “stress memory” has become increasingly critical, as it describes how plants retain information regarding past conditions to respond more effectively to similar future challenges (Ashapkin et al., 2020; Galviz et al., 2020). This phenomenon, often referred to as priming, entails an initial stress exposure leaving an imprint on the plant that influences its subsequent responses, allowing the organism to adapt and survive by anticipating future adversity (Haider et al., 2021). Such memory occurs when signals from an initial stress event are stored and later recalled, providing an integral component of plant resilience in a changing climate (Lämke and Bäurle, 2017). By enabling an enhanced or swifter response to subsequent stressors, this memory can significantly augment a plant’s overall tolerance (Crisp et al., 2016; Ramírez et al., 2015).

Epigenetic mechanisms, which do not involve DNA sequence alterations, are crucial for acquiring, retaining, and transmitting memory. Given the extensive chromatin reorganization during cell division, the transmission of parental memory likely requires specific mechanisms depending on the mode of reproduction. In sexually reproducing species, epigenetic information can be inherited through meiosis (Bird, 2007; Crisp et al., 2016). In clonal plants, epigenetic inheritance can occur through mitosis via stem cells in meristems (Gallusci et al., 2023). Interestingly, Latzel et al. (2016) found that clonal plants exhibit enhanced recovery from stress and improved performance under adverse conditions compared to non-clonal plants, suggesting a greater capacity to retain and use stress-induced molecular marks.

Memory formation is influenced by multiple factors, including differential sensitivity among plant organs, as well as the severity and duration of stress, genotype, and the developmental stage at which stress occurs (Cavatte et al., 2012). In wheat, for instance, priming has been shown to be more effective in alleviating heat and drought stress in susceptible cultivars than in tolerant ones, highlighting the genotype-specific nature of priming responses (Mendanha et al., 2020). In addition, the developmental stage plays a critical role in determining how stress memory is acquired and expressed, providing insight into the temporal dynamics of this process throughout the plant life cycle. Consequently, the same plant may exhibit distinct memory responses depending on the timing of stress exposure (Auler et al., 2021; Kron et al., 2008).

Recent findings suggest that the persistence of key metabolites following stress recovery may create a metabolite imprint, preparing plants for future adverse conditions (Schwachtje et al., 2019). In a GC-MS-based metabolic profiling study, Yadav et al. (2019) examined drought tolerance mechanisms in several wheat cultivars and found that primed plants under drought stress presented higher levels of glutamine, serine, methionine, lysine, and asparagine. Osmotic adjustment, a crucial mechanism for maintaining plant water status, is also closely associated with drought memory (Jacques et al., 2021). Several studies have reported that proline, a key osmolyte involved in drought tolerance, accumulates to higher levels in leaves following subsequent drought exposure, further supporting its role in drought stress memory (Qin et al., 2017; Marcos et al., 2018a).

Despite increasing evidence for stress memory in plants, little is known about how recurrent drought shapes physiological and metabolic responses in sugarcane or whether such responses persist across vegetative propagation. Herein, we evaluated how previous drought events affect subsequent responses in two sugarcane genotypes subjected to water deficit at distinct developmental stages, as well as the transmission of drought memory to vegetatively propagated clonal plants. We hypothesized that drought memory would be established when water deficit occurs during tillering, a developmental stage highly sensitive to water limitation (Machado et al., 2009). In contrast, drought imposed during the maturation may not promote memory acquisition. This hypothesis is consistent with evidence that stress occurring at later developmental stages can increase plant susceptibility to subsequent stress events, as reported in soybean exposed to drought during the reproductive stage (Kron et al., 2008).

## 2. Materials and methods

### 2.1 Plant material and growth conditions

Three-month-old sugarcane seedlings (*Saccharum* spp.) of IACSP95-5000 (high productivity) and IACCTC07-8008 (drought-tolerant) genotypes, obtained from the ProCana Breeding Program at the Agronomic Institute (IAC), Brazil, were planted in 20 L pots filled with dystrophic red latosol. The soil was fertilized with ammonium sulfate (750 mg kg^-1^), simple superphosphate (1350 mg kg^-1^), potassium chloride (400 mg kg^-1^), dolomitic limestone (600 mg kg^-1^), iron chelate (Fe 6%; 750 µg kg^-1^), and a micronutrient mixture (B 4.1%, Cu-EDTA 4.09%, Mn-EDTA 4.09%, Mo 0.92%, Ni 0.81%, Zn-EDTA 1.6%; 450 µg kg^-1^). The plants were irrigated daily and cultivated in a greenhouse with monitored temperature and relative humidity using a model HMP45C sensor (Campbell-Scientific, Logan UT, USA). Hourly data were recorded by a CR800 datalogger (Campbell-Scientific, Logan UT, USA) (Fig. S1).

#### 2.1.1 Phase I: inducing drought stress memory in the origin material

In the fifth month post-transplanting, the plants were randomly assigned to three experimental groups: well-watered (W), tillering (T) and maturation (M). Group W remained under daily irrigation throughout the experimental period, while groups T and M were subjected to three cycles of water deficit (D1, D2 and D3) during the tillering (group T, 5 months) and maturation (group M, 8 months) stages, respectively (Fig. 1a). Water deficit cycles during the maturation stage were imposed when the maturation index (MI) of the stalk reached 0.7, calculated as the ratio of soluble solids content in the apical part to that in the basal part (MI= °Brix apical/°Brix basal). The induction of drought memory in sugarcane plants was carried out according to Marcos et al. (2018b). Each water deficit cycle consisted of nine days of water withholding, followed by a six-day recovery period with daily irrigation (R). Soil moisture was monitored with Watermark 200SS sensors (Irrometer Co. Riverside CA, USA) (Fig. S1).

**Figure 1.**
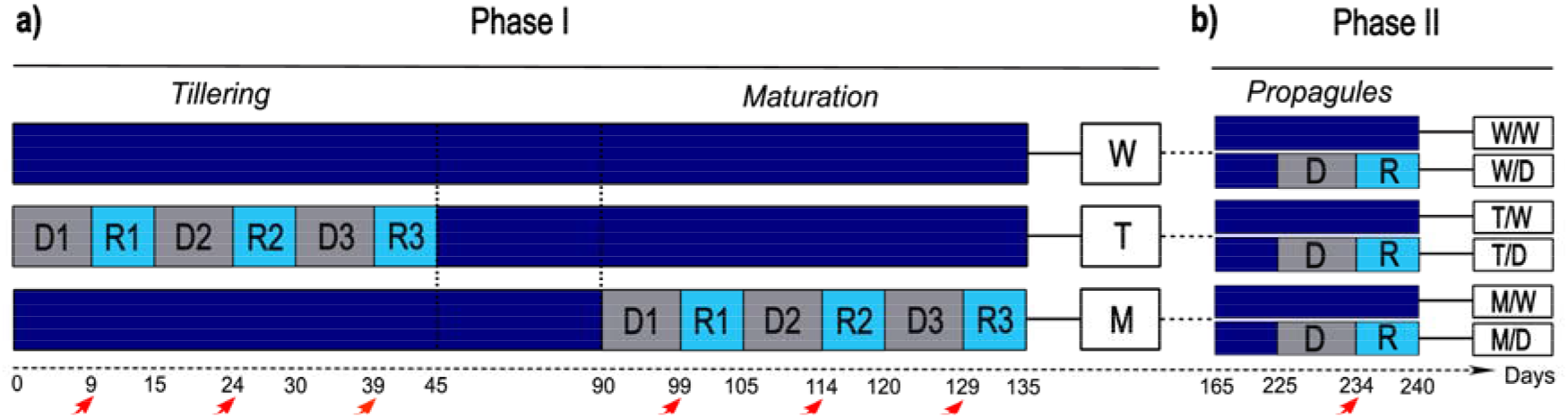
Schematic representation of drought cycles in two sugarcane genotypes (IACCTC07-8008 and IACSP95-5000) at two phenological stages (tillering and maturation) during phase I (origin material, in a) and phase II (propagules, in b). Shaded grey squares indicate drought cycles (D), while light blue squares represent recovery periods (R). Dark blue areas indicate plants well-watered. W= well-watered; T= plants subjected to three drought cycles and recovery during the tillering stage; M= plants subjected to three drought cycles and recovery during the maturation stage. W/W and W/D = well-watered and drought-stressed propagules, respectively, derived from well-watered plants; T/W and T/D = well-watered and drought-stressed propagules, respectively, derived from plants exposed to drought cycles at the tillering stage; M/W and M/D = well-watered and drought-stressed propagules, respectively, derived from plants exposed to drought cycles at the maturation stage. Red arrows indicate sampling at the end of each drought cycle (9^th^ day).

Leaves (+1, with a visible ligule) were collected on the day of maximum stress (9^th^ day of water withholding) of each water deficit cycle (red arrows, Fig. 1), immediately frozen in liquid nitrogen, and stored at −80 °C. Subsequently, setts containing a bud from all three groups of both genotypes were collected and planted in 500 mL pots containing a substrate of *Sphagnum* spp., rice straw, and perlite (7:2:1, Carolina Soil of Brazil, Vera Cruz, RS). The seedlings were maintained in a greenhouse under daily irrigation.

#### 2.1.2 Phase II: inducing drought stress in propagules

One month after planting, the vegetatively propagated seedlings, hereafter referred to as propagules, were transplanted into 12 L pots containing dystrophic red latosol. The pots were fertilized with urea (33.4 g pot^-1^, equivalent to 300 kg N ha^-1^), superphosphate (83.4 g pot^-1^, equivalent to 300 kg P_2_O_5_ ha^-1^), and potassium chloride (21.4 g pot^-1^, equivalent to 260 kg KCl ha^-1^) following the method described by Dias and Rossetto (2006). The propagules were maintained under conditions similar to those used for the original plants of phase I. After two months, they were divided into two water treatments: well-watered (W) and water deficit (D). In this manner, propagules derived from the three origin plant groups (W, T, and M) of both genotypes were subjected to two water regimes, resulting in six treatment combinations (W/W, W/D, T/W, T/D, M/W, and M/D) (Fig. 1b). The water deficit (D) treatment consisted of nine days of water withholding, followed by six days of recovery (R), after which the experiment was concluded. Soil moisture was monitored with Watermark 200SS sensors (Irrometer Co. Riverside CA, USA) (Fig. S1).

### 2.2 Leaf water status

Leaf water potential (Ψ_W_) was acquired with a pressure chamber (Model 1000, PMS Instrument Company, Albany OR, USA) on the day of maximum water deficit (9^th^ day). Measurements were taken on expanded leaves, collected around noon (11:30 am − 12:30 pm). At the same time, leaf relative water content (RWC) was determined from the fresh (FW), turgid (TW) and dry (DW) weights of leaf discs (20 mm), according to Jamaux et al. (1997):

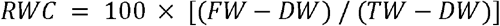

### 2.3 Leaf gas exchange and photochemical activity

Gas exchange and photochemical activity of the first fully expanded leaf with a visible ligule were measured throughout the experimental periods using an infrared gas analyzer (model LI-6400XT, LI-COR Inc., Lincoln NE, USA) with an attached modulated fluorometer (model 6400-40, LI-COR Inc., Lincoln NE, USA). Net CO_2_ assimilation (*A*_n_), stomatal conductance (*g*_s_), transpiration (E) and effective quantum efficiency of photosystem II (Φ_PSII_) were measured under PPFD of 2000 μmol m^−2^ s^−1^ and air CO_2_ concentration of 400 μmol mol^−1^. The instantaneous carboxylation efficiency (k_c_ = *A*_n_/C_i_) and intrinsic water use efficiency (WUE_i_ = *A*_n_/*g*_s_) were calculated according to Machado et al. (2009). The reductions in *A* and *g* were quantified by calculating the area under the curve over time for each drought cycle (drought/control), allowing the assessment of cumulative drought effects throughout the experimental period. By analyzing chlorophyll fluorescence, we estimated the maximum quantum efficiency of photosystem II (F_v_/F_m_) in leaves that were dark-adapted for 30 minutes. The measurements were performed between 10:30 and 12:30 h, as carried out by Pissolato et al. (2020).

### 2.4 Primary metabolism profile

Metabolite extraction and derivatization were conducted following the procedures outlined by Lisec et al. (2006) and Tohge and Fernie (2010). In brief, approximately 50 mg of lyophilized leaves were disrupted by shaking the cells in an Eppendorf tube with metal balls. The extraction was then carried out in 1.4 ml of methanol (LC–MS grade, Sigma-Aldrich®), stirring at 70 °C for 1 hour; 60 μl of ribitol (0.2 mg ml^-1^; Sigma-Aldrich®) was added as an internal standard to account for potential experimental errors during sample preparation and variations in GC–MS sensitivity (Lisec et al. 2006). After centrifugation at 11,000 *g* for 10 min, the supernatant was collected, and the polar and nonpolar phases were separated by adding 375 µL of chloroform (LC–MS grade, Sigma-Aldrich®) and 700 µL of deionized water. A new centrifugation (10,000 *g*, 15 min) was performed, and 1 ml of the polar phase (upper) was removed and dried using a vacuum lyophilizer. Subsequently, samples were derivatized by adding 40 µL of methoxyamine hydrochloride (Sigma-Aldrich®) in 20 mg/mL solution in pyridine (Sigma-Aldrich®) and incubated at 37 °C for 2 h. After derivatization, the samples were analyzed by gas chromatography coupled to mass spectrometry (GC-MS).

### 2.5 Plant growth analysis

The leaf and root dry mass were determined by drying the samples in an oven at 60 °C with forced air circulation until constant weight. The leaf area of each plant was assessed using a portable leaf area meter (model LI-3000, Li-Cor Inc., Lincoln NE, USA).

### 2.6 Data analysis

The data were subjected to Bayesian statistical analysis using the JASP software (https://jasp-stats.org/). In cases where significant differences were identified, the mean values were compared using the Bayes Factor (BF_10_). Our interpretation of the Bayes Factor as evidence for an alternative hypothesis (H_1_) was based on Miranda (2021): when 1 < BF_10_ < 3, there is weak support for H_1_; 3 < BF_10_ < 20 indicates positive support for H_1_; and BF_10_ > 20 indicates strong support for the alternative hypothesis.

Regarding the analysis of primary metabolism profile data, the relative abundance of metabolites was performed using the Xcalibur® 2.1 software (Thermo Fisher Scientific, https://www.thermofisher.com/). Each analyte peak was normalized to the peak intensity of the internal standard (ribitol). Pairwise differences between drought-treated plants and their respective well-watered controls were assessed for each metabolite using two-sided Welch’s t-tests. Log_2_ fold change (log_2_FC) was calculated as the log_2_-transformed ratio between mean metabolite abundance under drought and the corresponding well-watered control. Metabolites were considered differentially accumulated when p < 0.05 and |log_2_FC| > 1. These criteria were used for volcano plots, metabolic heatmaps, and supplementary tables. Principal component analysis (PCA) was performed in RBio software (Rbio 143, Viçosa MG, Brazil). PCA scores were visualized using the first two principal components. Shaded ellipses were generated using a multivariate t-distribution to visualize within-group dispersion.

## 3. Results

### 3.1. Leaf water status and gas exchange

In both genotypes, relative water content (RWC) remained high under well-watered conditions at both the tillering and maturation stages. In contrast, the first drought cycle significantly reduced RWC (Fig. S2a-f). At the tillering stage, both genotypes showed a greater RWC in D2 and D3 compared with D1 (Fig. S2a,d). During maturation, IACSP95-5000 showed a significant increase in RWC at D3 (Fig. S2b), whereas in IACCTC07-8008 a significant increase was already observed at D2 (Fig. S2e). In Phase II, propagules under drought and derived from drought-stressed plants (T/D and M/D) showed higher RWC than W/D in both genotypes (Fig. S2c,f); however, differences were not statistically significant in IACSP95-5000 (Fig. S2c). Regarding leaf water potential (Ψ_w_), no significant differences among drought cycles in IACSP95-5000 either during Phase I or in propagules (Fig. S2g–i). In contrast, IACCTC07-8008 showed less negative Ψ_w_ values in D2 and D3 compared with D1 during maturation (Fig. S2k). In Phase II, only M/D presented lower Ψ_w_ compared with D1, while the remaining treatments did not differ (Fig. S2l).

In IACSP95-5000, recurrent drought cycles caused pronounced reductions in *A*_n_, *g*_s_, E, and k_c_ during tillering, with all parameters decreasing sharply as water deficit progressed and reaching their lowest values in the maximum drought intensity (9^th^ day of each cycle) (Fig. S3a,d,g,j). Following rehydration, gas exchange parameters recovered progressively, with a noticeably faster recovery during the second (R2) and third (R3) recovery periods compared with the first period (R1). At maturation, a similar cyclic response was observed, characterized by marked declines in all evaluated parameters during drought periods followed by recovery after rehydration (Fig. S3b,e,h,k). Well-watered plants maintained comparatively stable physiological performance throughout both developmental stages, despite some temporal fluctuations.

In IACCTC07-8008, *A* declined sharply during each drought cycle, reaching minimum values at maximum water deficit, with similar dynamics observed for *g*_s_ and E, and a pronounced decrease in k_c_ (Fig. S4a,d,g,j). At maturation, a comparable oscillatory pattern was observed, characterized by reductions during drought periods and recovery upon rehydration (Fig. S4b,e,h,k). Well-watered plants maintained relatively stable physiological performance throughout the experimental period.

In Phase II, propagules under drought of both genotypes showed reductions in *A*_n_, *g*_s_, E, and k_c_ relative to well-watered plants, with responses influenced by parental origin. In IACSP95-5000, propagules under drought derived from well-watered plants (W/D) exhibited the lowest *A*_n_ and k_c_ values at maximum water deficit (Fig. S3c,l), whereas propagules originating from plants previously exposed to drought at tillering (T/D) or maturation (M/D) showed a faster recovery of *A*_n_, *g*_s_, E, and k_c_ following rewatering (Fig. S3c,f,i,l). A similar pattern was observed in IACCTC07-8008, where propagules under drought reached minimum values at maximum stress (Fig. S4c,f,i,l). Although no differences were detected among W/D, T/D, and M/D at the end of the stress period, propagules derived from drought-stressed plants recovered more rapidly following rewatering.

The integrated analysis of drought-induced reductions revealed significant differences among drought cycles and propagules in both genotypes (Fig. 2). In IACSP95-5000, cumulative reductions in *A*_n_ were greatest during D1 at both tillering and maturation, with progressively smaller effects in subsequent cycles, particularly D2 (Fig. 2a,b). A similar trend was observed in IACCTC07-8008 at tillering, while at maturation D1 and D2 showed greater effects than D3 (Fig. 2d,e). In Phase II, propagules under drought derived from well-watered plants (W/D) exhibited greater cumulative reductions in *A*_n_ than those derived from drought-stressed plants (T/D and M/D) in both genotypes (Fig. 2c,f). A similar pattern was observed for *g*_s_ in IACSP95-5000 (Fig. 2g–i). In IACCTC07-8008, *g*_s_ reductions followed the same cycle-dependent pattern, with the largest effects in D1 at tillering and reduced impact in later cycles, while at maturation D1 induced greater reductions than D3 (Fig. 2j,k). Consistently, propagules derived from drought-stressed plants showed smaller cumulative reductions compared with the W/D group (Fig. 2l).

**Figure 2.**
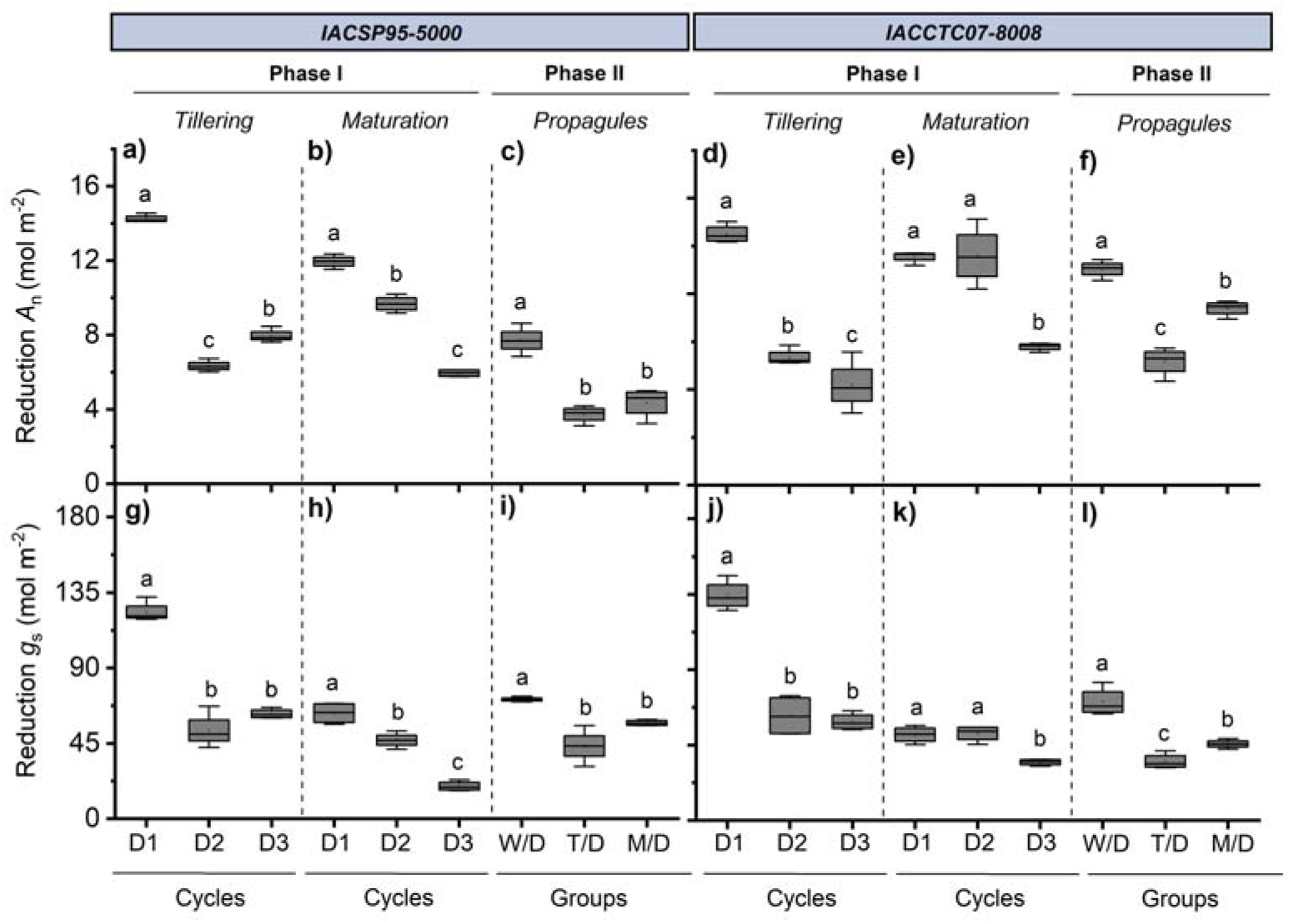
Reduction in net CO_2_ assimilation (Reduction *A*, in a-f) and stomatal conductance (Reduction *g*, in g-l) integrated across drought cycles in phase I and phase II in two sugarcane genotypes (IACSP95-5000 and IACCTC07-8008). Boxplots represent the median (centre line), interquartile range (box limits), and data dispersion (whiskers). Different letters indicate significant differences among cycles or groups (BF_10_ > 3; n = 4).

Intrinsic water use efficiency (WUE_i_), measured at maximum water deficit, varied among drought cycles and propagule groups in both genotypes (Fig. 3). In Phase I at the tillering stage, IACSP95-5000 showed higher WUE_i_ in D3 than in D1 and D2, whereas no differences were observed among well-watered plants (Fig. 3a). At maturation, WUE_i_ was higher in D3 compared with D1 (Fig. 3b). In IACCTC07-8008, WUE_i_ at tillering was lowest in D1 and increased in subsequent cycles, reaching higher values in D3 (Fig. 3d), while at maturation it was lower in D1 than in D2 and D3, with no differences among well-watered cycles (Fig. 3e). In Phase II, WUE_i_ was influenced by parental history in both genotypes. In IACSP95-5000, well-watered propagules from drought-stressed plants (T/W and M/W) exhibited higher WUE_i_ than W/W (Fig. 3c). Under drought, propagules derived from drought-stressed plants (T/D and M/D) also showed higher WUE_i_ than those from well-watered parents (W/D), with the highest values observed in M/D (Fig. 3). A similar pattern was observed in IACCTC07-8008, where WUE_i_ was higher in T/W and M/W than in W/W under well-watered conditions, and in T/D and M/D than in W/D under drought (Fig. 3f).

**Figure 3.**
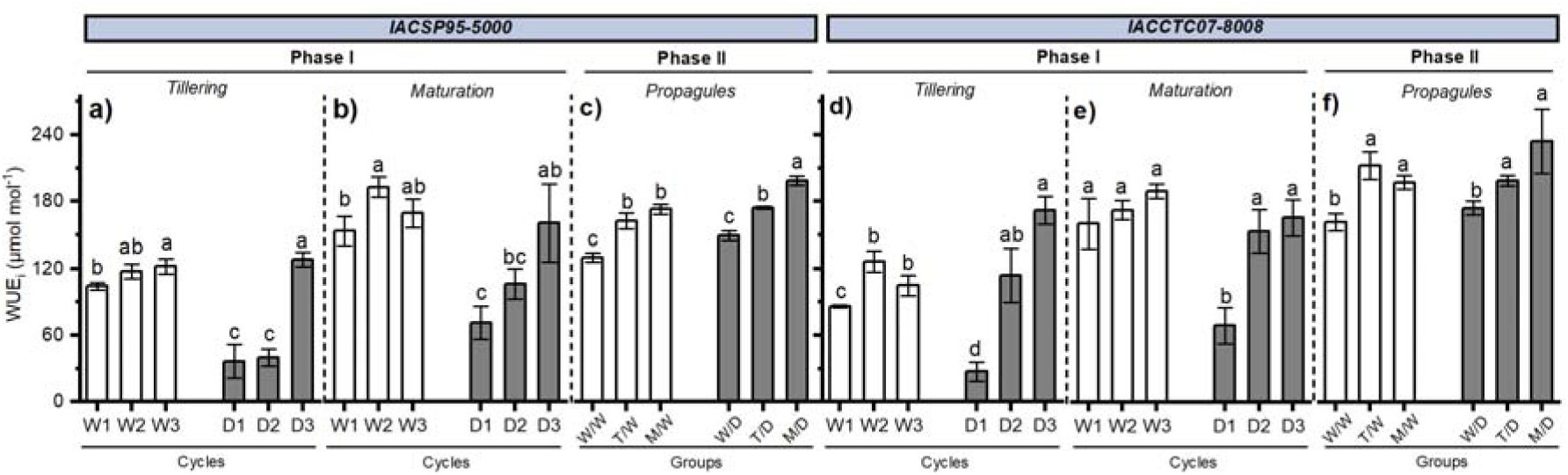
Intrinsic water use efficiency (WUE_i_) of two sugarcane genotypes (IACSP95-5000 and IACCTC07-8008) subjected to drought cycles (D) in phase I and phase II or kept well-watered (W). In phase I, plants were subjected to three drought cycles at the tillering or maturation stage, while in phase II, the propagules derived from well-watered plants (W/) or from plants previously exposed to drought at tillering (T/) or maturation (M/) and subsequently subjected to either drought (/D) or kept well-hydrated (/W). The data were collected during the maximum water deficit (9^th^ day). Bars represent means (n=4) ± SD. Different letters indicate significant differences among cycles or groups (BF_10_ > 3).

Maximum quantum efficiency of PSII (F_v_/F_m_) and effective quantum yield of PSII (Φ_PSII_) were measured at maximum water deficit across three drought cycles during tillering and maturation in both genotypes. At tillering, F_v_/F_m_ remained stable across well-watered cycles in both genotypes. Under drought, however, F_v_/F_m_ decreased significantly relative to well-watered plants (Fig. S5a-d). In IACSP95-5000, D1 resulted in lower F /F compared with D2 and D3 (Fig. S5a). In IACCTC07-8008, drought also reduced F_v_/F_m_, with D1 showing the lowest values and subsequent cycles exhibiting higher values, indicating differences among cycles under water deficit (Fig. S5c). At maturation, F_v_/F_m_ remained stable among well-watered cycles in both genotypes. No significant differences were observed among the groups in Phase II (data not shown).

At the tillering stage, the Φ_PSII_ did not differ among well-watered plants in both genotypes (Fig. S5e,g). Under drought, Φ_PSII_ declined during D1 and increased in subsequent cycles in IACSP95-5000 (Fig. S5e), whereas in IACCTC07-8008 this increase was only evident in D3 (Fig. S5g). At maturation, IACSP95-5000 showed higher Φ_PSII_ in D3 compared with D1 and D2 (Fig. S5f), and a similar pattern was observed in IACCTC07-8008, where D3 exhibited higher values than D1 (Fig. S5h). No significant differences were detected among groups in Phase II (data not shown).

### 3.2 Metabolite trends under recurrent drought

Volcano plot analyses revealed differential regulation of primary metabolites in response to drought across developmental stages (Phase I; Fig. S6) and propagule groups (Phase II; Fig. S7) in both genotypes. In total, we identified 38 metabolites modulated by water deficit cycles in leaves, including mainly amino acids, organic acids, sugars, and their derivatives. Detailed metabolite identities, fold changes, and significance values are presented in Tables S1, S2 and S3.

In Phase I, drought induced consistent metabolic reprogramming in both genotypes, characterized predominantly by the accumulation of amino acids and sugars. At the tillering stage in IACSP95-5000, drought promoted the upregulation of several amino acids, including valine, leucine, isoleucine, glycine, serine, proline, and phenylalanine, along with sugars such as glucose and fructose (Fig. S6a–c; Table S1). A similar pattern was observed in IACCTC07-8008, where drought enhanced the accumulation of branched-chain amino acids and osmoprotective compounds such as glycine and proline across cycles (Fig. S6d–f). In addition, drought modulated metabolites associated with central metabolism, including organic acids such as malate, fumarate, and glutarate, with responses varying by cycle and developmental stage (Tables S1, S2). At maturation, both genotypes showed sustained increases in amino acids, particularly proline, serine, threonine, methionine, and phenylalanine, with more pronounced accumulation during later drought cycles (Fig. S6g–l). Drought cycles were consistently associated with a predominance of upregulated metabolites in both genotypes and developmental stages.

In Phase II (Fig. S7), metabolite regulatory patterns were evaluated in drought-stressed propagules compared with their respective well-watered controls. In IACSP95-5000, propagules subjected to drought and derived from well-watered plants (W/D vs. W/W; Fig. S7a) exhibited six upregulated metabolites (butanoate, glutarate, ornithine, fructose and glucose) and no downregulated metabolites. Propagules subjected to drought and derived from drought-stressed plants at tillering (T/D vs. T/W; Fig. S7b) showed 16 upregulated metabolites including valine, leucine, isoleucine, serine, and phenylalanine, as well as sugars such as fructose and glucose (Table S3), and no downregulated metabolites. We observed the same response for propagules subjected to drought and derived from drought-stressed plants at the maturation stage (M/D vs. M/W; Fig. S7c)

In IACCTC07-8008, propagules subjected to drought and derived from well-watered plants (W/D vs. W/W; Fig. S7d) showed 14 upregulated (including valine, isoleucine, glycine, serine, threonine and β-alanine) and one downregulated metabolite (glutarate) (Table S3). By contrast, propagules subjected to drought and derived from drought-stressed plants at tillering (T/D vs. T/W; Fig. S7e) exhibited 24 upregulated and 3 downregulated metabolites (malate, glutarate and sucrose). Finally, propagules subjected to drought and derived from drought-stressed plants at maturation stage (M/D vs. M/W; Fig. S7f) revealed 21 upregulated (including valine, leucine, isoleucine, glycine and proline) and three downregulated metabolites (urea, malate and glutarate).

Principal component analysis (PCA) revealed clear separation among drought cycles and propagule groups in both genotypes (Fig. 4). In Phase I, at the tillering stage, IACSP95-5000 showed separation between drought-treated and well-watered samples mainly along PC1, with D1 and D2 occupying more distinct positions and D3 showing a more intermediate distribution (Fig. 4a). A similar pattern was observed in IACCTC07-8008, in which drought-treated samples were separated from well-watered plants primarily along PC1 (Fig. 4c). At the maturation stage, both genotypes also showed clear discrimination between drought-treated and well-watered samples, while PC2 contributed to the separation among drought cycles, especially for D1 in IACSP95-5000 and D3 in IACCTC07-8008 (Fig. 4b,d). PC1 explained most of the variance and primarily distinguished drought-treated from well-watered plants, whereas PC2 accounted for additional variation among drought cycles.

**Figure 4.**
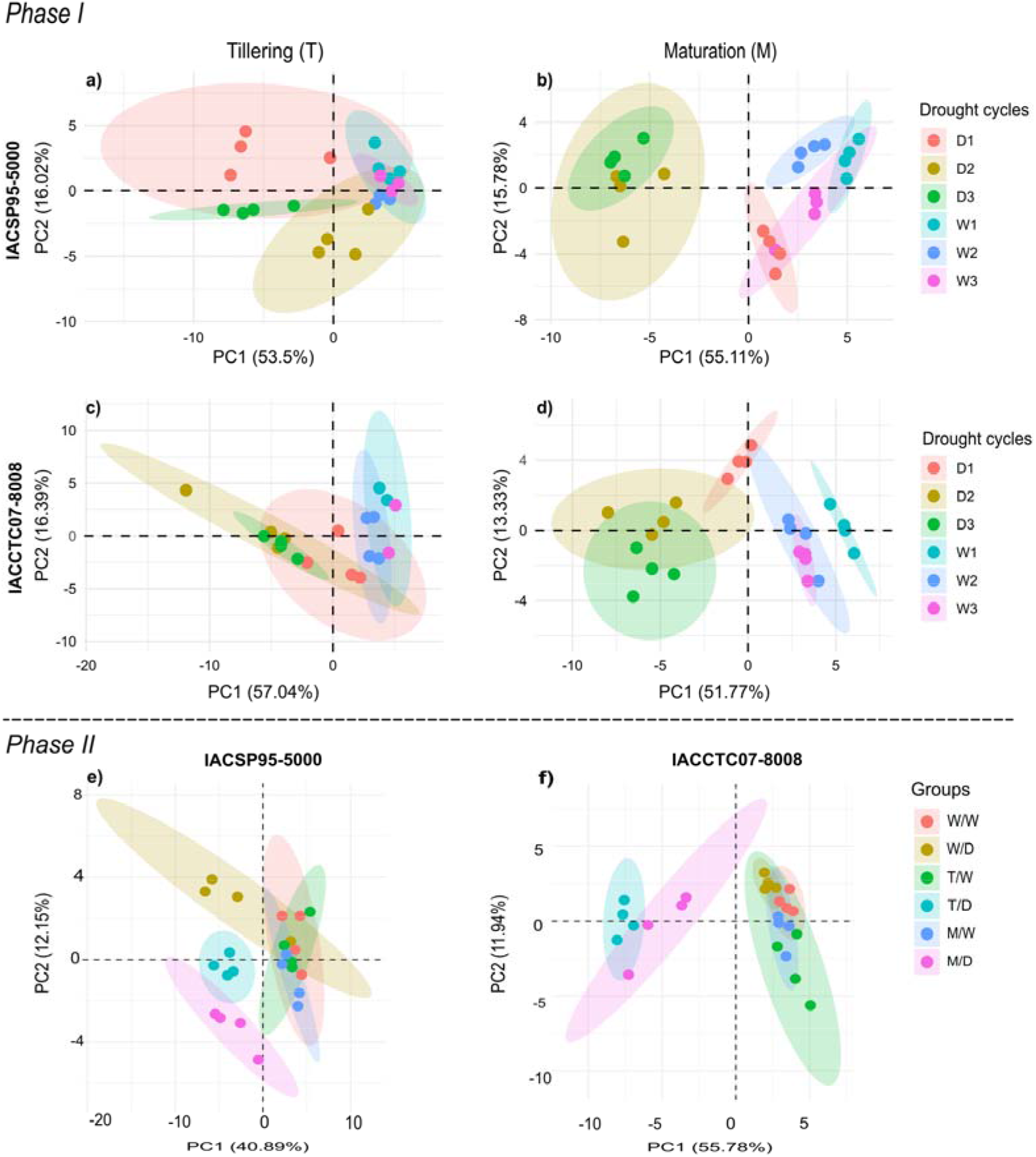
Principal component analysis (PCA) of leaf metabolites from two sugarcane genotypes IACSP95-5000 and IACCTC07-8008. Phase I depicts plants subjected to three drought cycles (D) at the tillering (T) or maturation stage (M) or kept well-watered (W) (in a-d). Phase II shows sugarcane propagules derived from well-watered plants or from plants previously exposed to drought cycles at the tillering or maturation stage, which were subsequently either subjected to drought (/D) or maintained under well-watered conditions (/W) (in e-f). Points represent biological replicates for each group. Shaded ellipses indicate the dispersion of biological replicates within each treatment group. Dashed lines represent zero reference axes. The PC1 and PC2 axes display the percentage of variance explained by each principal component.

In Phase II, separation among propagule groups was observed in both genotypes. In IACSP95-5000 (Fig. 4e), groups derived from well-watered plants and those derived from drought-stressed plants at tillering or maturation stages showed partial clustering, with separation mainly along PC1. In IACCTC07-8008 (Fig. 4f), a clearer separation was observed, particularly along PC1, distinguishing propagules derived from drought-stressed plants from those derived from well-watered plants. M/D samples were positioned on the negative side of PC1, while T/W and M/W clustered on the positive side, and W/D samples showed an intermediate distribution.

Volcano plot analysis provided an overview of the magnitude and direction of the metabolic response to drought across cycles, developmental stages, and genotypes (Figs. S6, S7). During Phase I, upregulation predominated in most comparisons, although the number of significantly regulated metabolites varied among drought cycles. At tillering, IACSP95-5000 showed 20 upregulated and no downregulated metabolites in D1, 7 upregulated and none downregulated in D2, and 20 upregulated and 2 downregulated in D3. In IACCTC07-8008, D1 resulted in 11 upregulated and 5 downregulated metabolites, D2 in 14 upregulated and 3 downregulated metabolites, and D3 in 19 upregulated and 4 downregulated metabolites. At maturation, IACSP95-5000 exhibited 16 upregulated and 4 downregulated metabolites in D1, 21 upregulated and 2 downregulated in D2, and 20 upregulated and 2 downregulated in D3. In IACCTC07-8008, 14 metabolites were upregulated and 6 downregulated in D1, 15 were upregulated and 3 downregulated in D2, and 17 were upregulated and 2 downregulated in D3 (Fig. S6).

During Phase II, drought also resulted predominantly in metabolite upregulation, with the magnitude of the response varying according to the drought history of the source plants. In IACSP95-5000, propagules derived from well-watered plants (W/D) showed 6 upregulated and no downregulated metabolites, whereas propagules derived from plants previously exposed to drought at tillering (T/D) and maturation (M/D) each showed 16 upregulated metabolites and no downregulated metabolites (Fig. S7a-c). In IACCTC07-8008, W/D propagules exhibited 14 upregulated and 1 downregulated metabolite, while T/D propagules showed 24 upregulated and 3 downregulated metabolites and M/D propagules showed 21 upregulated and 3 downregulated metabolites (Fig. S7d-f). Overall, propagules originating from drought-exposed plants displayed a greater number of significantly regulated metabolites than those derived from well-watered plants, particularly in IACCTC07-8008.

Metabolic maps were then used to identify the metabolic pathways underlying these global changes (Figs. 5, 6). During Phase I, drought prominently affected amino acid metabolism, carbohydrate metabolism, the TCA cycle, and the shikimate pathway in both genotypes (Fig. 5). A common response across developmental stages and drought cycles was the accumulation of several amino acids, particularly branched-chain amino acids, threonine, β-alanine, glycine, serine, proline, and phenylalanine. In parallel, central carbon metabolism showed a contrasting pattern, with several TCA cycle intermediates, including malate and fumarate, generally displaying negative log_2_ fold changes, whereas glucose and fructose were predominantly positively regulated.

**Figure 5.**
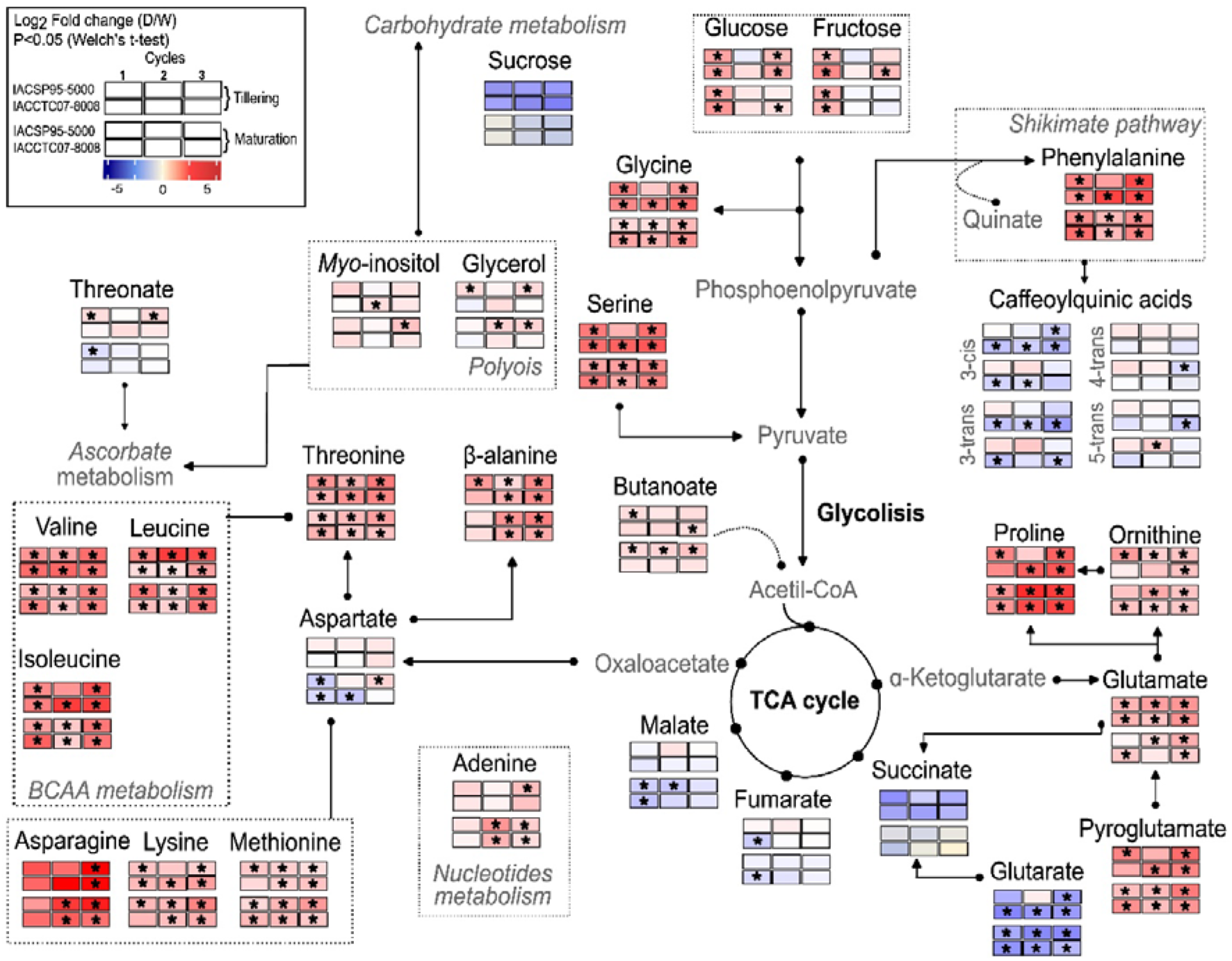
Metabolic map showing the log fold change (D/W) of leaf metabolites significantly affected by drought in two sugarcane genotypes (IACSP95-5000 and IACCTC07-8008) subjected to three water-deficit cycles at the tillering or maturation stage. Differential metabolite accumulation was assessed using two-sided Welch’s t-tests. Only metabolites meeting both p < 0.05 and |log_2_FC| > 1 were included. Positive log_2_FC values indicate increased metabolite abundance under drought, whereas negative values indicate decreased abundance relative to the corresponding well-watered control. Asterisks indicate metabolites meeting the statistical significance criterion (p < 0.05, Welch’s t-test).

Some responses were dependent on genotype, developmental stage, or drought cycle. In IACSP95-5000, branched-chain amino acids showed particularly consistent positive regulation throughout tillering and maturation, whereas aspartate was negatively regulated during the first drought cycle at maturation (Fig. 5). Sucrose and polyols, including *myo*-inositol and glycerol, exhibited more variable responses across cycles and genotypes. Metabolites associated with the shikimate pathway also showed coordinated regulation, with phenylalanine predominantly increasing under drought while caffeoyl shikimic acid showed mainly negative regulation (Fig. 5).

In Phase II, metabolic pathway analysis revealed that drought responses in the propagules were strongly influenced by the drought history of the source plants (Fig. 6). Propagules derived from plants previously exposed to drought at tillering or maturation generally showed stronger regulation of amino acid metabolism than those derived from well-watered plants. Branched-chain amino acids, threonine, β-alanine, asparagine, proline, glutamate, and pyroglutamate were predominantly positively regulated, together with glucose and fructose. In contrast, malate and fumarate remained negatively regulated, indicating a recurrent drought-associated response of the TCA cycle across generations. Sucrose, succinate, glutarate, *myo*-inositol, and glycerol showed more variable responses according to genotype and parental drought history (Fig. 6).

**Figure 6.**
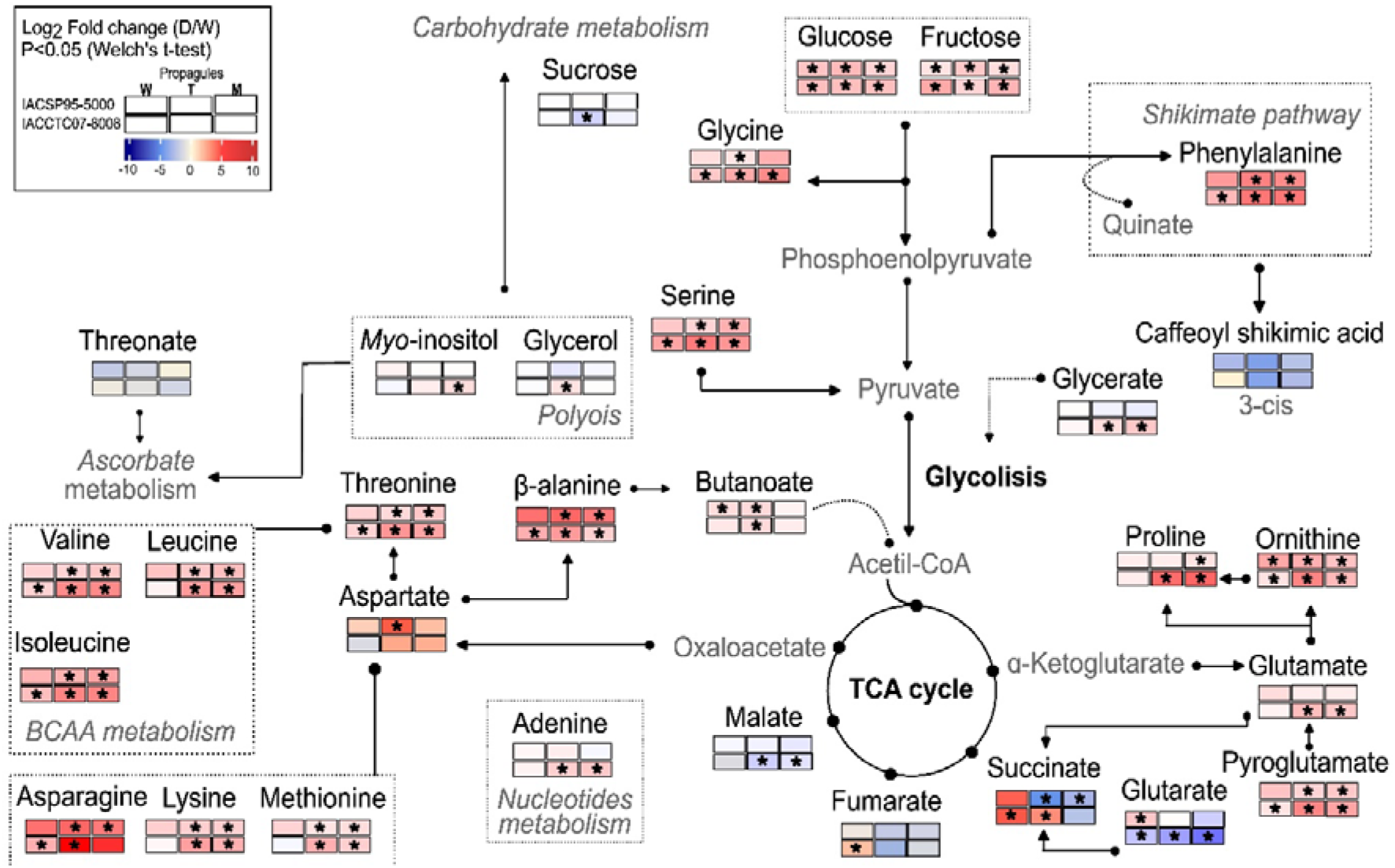
Metabolic map showing the log fold change (D/W) of metabolites significantly affected by drought in propagules of IACSP95-5000 and IACCTC07-8008 derived from well-watered plants (W/) or plants previously exposed to drought during tillering (T/) or maturation (M/). Propagules were subsequently subjected to drought (/D) or maintained under well-watered conditions (/W). Differential metabolite accumulation was assessed using two-sided Welch’s t-tests. Only metabolites meeting both p < 0.05 and |log_2_FC| > 1 were included. Asterisks indicate metabolites meeting the statistical significance criterion (p < 0.05, Welch’s t-test).

### 3.3 Plant growth

For both genotypes, shoot dry mass (SDM) in Phase I tended to be higher in drought-stressed plants at the tillering stage, although no significant differences were observed among groups (Fig. 7a,c). Both genotypes showed greater root dry mass (RDM) in both T and M stages compared with W (Fig. 7e,g). Leaf area of IACSP95-5000 was greater in T compared with W and M, but no differences were found in cv. IACCTC07-8008 (Fig. 7i,k).

**Figure 7.**
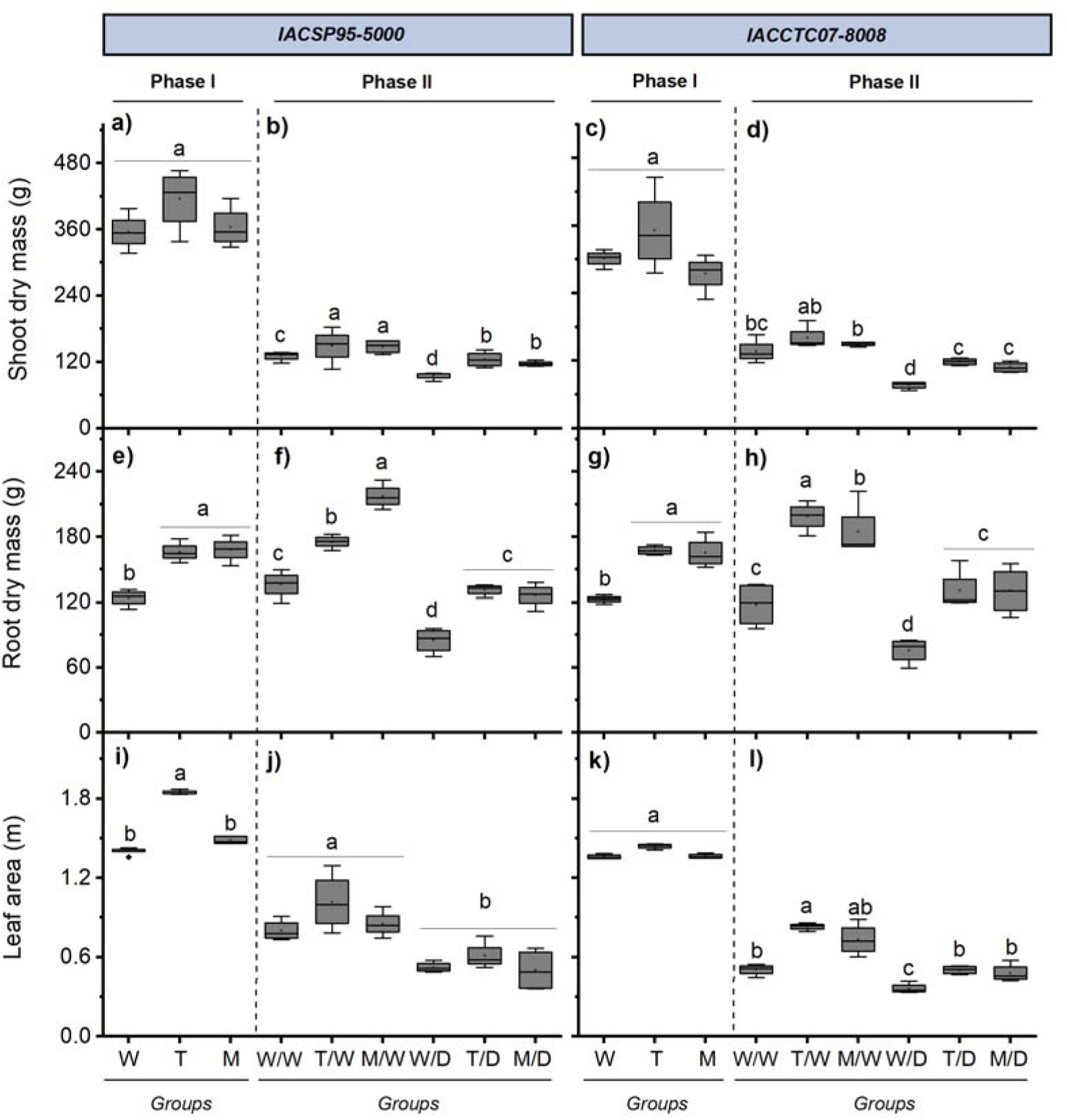
Shoot dry mass (a-d), root dry mass (e-h), and leaf area (i-l) in two sugarcane genotypes IACSP95-5000 and IACCTC07-8008. Phase I depicts plants subjected to three drought cycles at the tillering (T) or maturation stage (M) or kept well-watered (W). Phase II shows sugarcane propagules derived from well-watered plants (W/) or from plants previously exposed to drought cycles at the tillering (T/) or maturation stage (M/), which were subsequently either subjected to drought (/D) or maintained under well-watered conditions (/W). Boxplots represent the median (centre line), interquartile range (box limits), and data dispersion (whiskers). Different letters indicate significant differences among groups within each phase (BF_10_ > 3; n = 4).

In Phase II, SDM varied among propagule groups. Under well-watered conditions, the group T/W and M/W from IACSP95-5000 showed higher SDM than W/W (Fig. 7b). A similar pattern was observed under drought, where T/D and M/D exhibited greater SDM than W/D (Fig. 7b). In contrast, no significant differences were observed among propagule groups of IACCTC07-8008 under well-watered conditions; however, under drought, propagules derived from stressed plants (T/D and M/D) showed significantly higher SDM than W/D (Fig. 7d).

In IACSP95-5000, T/W exhibited the highest RDM among propagule groups, differing from W/W and M/W (Fig. 7f). Under drought, RDM decreased across all groups, with W/D showing the lowest values compared with T/D and M/D (Fig. 7f). In IACCTC07-8008, propagules derived from stressed plants (T/W and M/W) showed higher RDM than W/W under well-watered conditions, and a similar pattern was observed under drought, where T/D and M/D exhibited greater RDM than W/D (Fig. 7h). All propagule groups of IACSP95-5000 showed reduced leaf area under drought compared with irrigated conditions, with no differences among groups (Fig. 7j). In contrast, in IACCTC07-8008, T/W exhibited significantly greater leaf area than W/W, and under drought, both T/D and M/D also showed higher leaf area compared with W/D (Fig. 7i).

## 4. Discussion

### 4.1 Recurrent drought redefines the trajectory of the stress response in sugarcane

The first drought cycle (D1) imposed strong physiological limitations in both genotypes and developmental stages, as reflected by reductions in RWC, gas exchange, and photochemical parameters. In contrast, subsequent drought cycles (D2 and D3) resulted in attenuated reductions in *A*_n_ and *g*_s_, increased WUE_i_, and partial maintenance of photochemical performance. Recovery of photosynthetic traits also became faster during R2 and R3, indicating that the initial drought event acted as a conditioning stimulus. Together, these results demonstrate that recurrent drought attenuated reductions in *A*_n_ and accelerated post-stress recovery.

This response pattern was reinforced in Phase II. Under drought conditions, propagules derived from drought-stressed plants (T/D and M/D) exhibited significantly lower integrated reductions in *A*_n_ and *g*_s_ than propagules derived from well-watered plants (W/D) in both genotypes (Fig. 2c,f,i,l), indicating mitigation of cumulative physiological impact. Furthermore, propagules from drought-stressed plants restored *A*, *g* and k_c_ faster during rehydration than those from well-watered plants (Figs. S3-S4c,f,i,l), demonstrating that stress history enhanced both resistance to functional decline and recovery efficiency. Similar responses have been reported in sugarcane clones derived from drought-stressed plants, which showed faster recovery of photosynthetic parameters and higher k_c_ after rehydration (Marcos et al., 2018b). Also, unfertilized rice plants subjected to recurrent drought with a short recovery interval exhibited higher *A*_n_, WUE, and k_c_ compared with both unprimed plants within the same treatment group and plants grown under control conditions (Galviz et al., 2025).

Stress memory does not necessarily prevent initial physiological limitation; rather, it enhances the efficiency and speed of post-stress recovery. The faster recovery of *A*_n_ and k_c_ observed here defines a recovery-based resilience phenotype, consistent with frameworks proposing that memory frequently manifests through improved recovery kinetics rather than reduced peak impairment (Fàbregas and Fernie, 2019; Walter et al., 2011). The recovery phase itself appears central to memory consolidation. Following initial stress exposure, retained information may place plants in a permissive physiological state, enabling faster and more coordinated responses to subsequent stress events. This retention may involve accumulation of stress-related proteins in inactive forms (Virlouvet et al., 2018), persistence of specific metabolites and phytohormones, and epigenetics modifications including DNA methylation, histone modification, or chromatin remodeling (Hilker et al., 2015).

The progressive increase in WUE_i_, particularly during tillering for both genotypes (Fig. 3a,d), reflects enhanced integration between CO_2_ assimilation and stomatal conductance. Such improvement suggests refinement of guard cell responsiveness and potential recalibration of ABA-mediated signaling pathways (Virlouvet and Fromm, 2015; Brodribb et al., 2019). Similar memory-associated adjustments have been described in coffee, where prior drought exposure optimized photosynthetic performance and resource allocation, resulting in improved water use efficiency under subsequent stress (Menezes-Silva et al., 2017).

Recurrent drought was also associated with enhanced photochemical performance during drought stress cycles. F /F and Φ_PSII_ increased during D3 compared with D1 in both genotypes and at both developmental stages (Fig. S5), indicating improved maintenance of PSII functionality under repeated drought cycles. Importantly, this improvement coincided with higher RWC observed in later drought cycles (Fig. S2), suggesting that enhanced tissue hydration contributed to the preservation of photochemical integrity. Maintenance of cellular water status is known to stabilize thylakoid membranes, reduce excessive excitation pressure, and limit reactive oxygen species formation under drought conditions (Flexas et al., 2014). The increase in Φ_PSII_ during D3 (Fig. S5e-h) indicates more efficient allocation of absorbed light energy to photochemistry, which may have supported the improved photosynthetic performance observed under recurrent drought. These responses demonstrate that recurrent drought reinforced the stability of the photosynthetic apparatus rather than intensifying photoinhibitory damage. Interestingly, while this photochemical reinforcement was evident in plants directly exposed to recurrent drought (Phase I), similar differences were not detected in vegetatively derived propagules under subsequent stress (data not shown). This indicates that improvements in recovery kinetics were retained across propagation, whereas adjustments in photochemical efficiency may depend more directly on repeated stress exposure. Collectively, these findings suggest that recurrent drought improved the coordination between water status and photochemical performance, contributing to the establishment of a physiologically stabilized, memory-associated state.

### 4.2 Metabolic reprogramming underlies memory

Recurrent drought induced sustained reorganization of primary metabolism in sugarcane leaves, particularly affecting amino acid and central carbon metabolism. Branched-chain amino acids (valine, leucine, and isoleucine) remained consistently upregulated during D2 and D3, indicating that metabolic responses were reinforced rather than attenuated across stress cycles (Tables S1–S3). Similar metabolic adjustments have been reported in drought-stressed maize and wheat, where amino acid accumulation and carbon reallocation were associated with stress acclimation and metabolic preparedness (Li et al., 2024; Itam et al., 2020). The persistence of these metabolic configurations supports the concept of “metabolic imprinting,” in which prior stress exposure establishes a preconditioned metabolic state that influences subsequent drought responses (Schwachtje et al., 2019). Within this broader metabolic reprogramming, the accumulation of specific amino acids with protective roles becomes particularly relevant. Notably, the increased levels of proline observed during D3 for both genotypes and developmental stages reinforce its function as a key osmoprotectant under drought stress. Osmotic adjustment for the maintenance of plant water status has been implicated as a central component of drought stress memory (Jacques et al., 2021), and proline plays a major role in this process due to its function as an osmolyte. Menezes-Silva et al. (2017) demonstrated that coffee plants exposed to recurrent drought events exhibit adaptive responses to subsequent stress, driven by the expression of trainable genes linked to drought tolerance and supported by extensive metabolic reprogramming. In this context, transcriptional memory of Δ¹-pyrroline-5-carboxylate synthetase 1 (P5CS1), a key enzyme in proline biosynthesis, has been identified as a critical component of drought stress memory in rice, reinforcing the link between proline metabolism and priming mechanisms.

Consistent with this interpretation, increased proline accumulation under recurrent drought has been reported in multiple species, including *Dipteryx alata* L. (Alves et al., 2020) and peanut plants (Qin et al., 2017), suggesting that enhanced proline levels may represent a conserved adaptive response. However, contrasting patterns have also been observed. Studies in sugar beet (Leufen et al., 2016) and soybean (Nguyen et al., 2020) reported reduced proline accumulation during subsequent drought events compared to the initial stress, indicating that proline dynamics may be species-dependent and influenced by the intensity and frequency of stress exposure. The sustained accumulation of proline observed in this study supports its role not only as an osmoprotector but also as a component of metabolic imprinting associated with drought stress memory, potentially enhancing the plant’s capacity to maintain cellular homeostasis under recurrent stress conditions. Similar evidence has been reported in sugarcane, where propagules derived from plants previously exposed to water deficit exhibited higher proline accumulation, improved photosynthetic performance, and enhanced drought tolerance compared with propagules from non-stressed plants (Marcos et al., 2018b).

The accumulation of asparagine, particularly in D3 compared to D1 (Fig. 5), indicates enhanced nitrogen remobilization and redistribution during recurrent drought cycles. Consistent with this interpretation, asparagine accumulation under water deficit has been associated with adjustments in nitrogen metabolism and source–sink dynamics, as well as with the maintenance of metabolic activity and improved stress tolerance in plants (Rubia et al., 2025; Akin and Kaya, 2024). In parallel, branched-chain amino acids, including valine, leucine and isoleucine, were also upregulated, supporting their role as alternative respiratory substrates and contributors to metabolic flexibility under stress (Araújo et al., 2011; Pires et al., 2016). Consistently, wheat plants previously exposed to drought showed increased levels of branched-chain amino acids upon re-exposure to water deficit (Li et al., 2024), supporting the interpretation that amino acid accumulation represents a coordinated metabolic strategy associated with stress preparedness. In coffee plants exposed to repeated drought, increases in glycine, isoleucine, phenylalanine, glycerol and *myo*-inositol have been associated with the establishment of an “alert metabolic state” (Menezes-Silva et al., 2017), consistent with the broader metabolic adjustments and reinforced metabolite pools observed here.

Concomitantly, modulation of TCA cycle intermediates, including reductions in malate and fumarate in later cycles, indicates reconfiguration of respiratory metabolism under restricted carbon assimilation (Fàbregas and Fernie, 2019). Such coordinated shifts in glycolysis- and TCA-related pathways have similarly been reported in wheat under drought, where central carbon metabolism was tightly integrated with amino acid biosynthesis and stress-responsive regulation (Li et al., 2025). Importantly, this metabolic restructuring intensified across cycles, as reflected by the predominance of upregulated metabolites during D2 and D3 relative to D1, suggesting amplification rather than attenuation of metabolic responsiveness. A comparable progression was described in wheat subjected to progressive drought, where metabolomic transitions followed defined physiological thresholds and involved coordinated regulation of amino acid, nucleotide and TCA-cycle metabolism (Itam et al., 2020). In that study, severe drought induced strong accumulation of branched-chain amino acids and purine-related metabolites, reflecting nitrogen recycling and metabolic adjustment to carbon limitation. In contrast to progressive drought systems, however, our results indicate that cyclic drought exposure did not culminate in metabolic collapse but instead established a reinforced and anticipatory metabolic configuration. The persistence of elevated amino acid pools and shikimate pathway activation (Figs. 5, 6) suggests that recurrent drought reshaped central carbon–nitrogen balance in a manner consistent with metabolic imprinting, particularly evident in propagules derived from previously stressed plants.

The reduction of malate and fumarate in later cycles indicates an adjustment in carbon flux under restricted assimilation, potentially redirecting carbon skeletons toward protective biosynthesis. Notably, activation of the shikimate pathway, reflected by sustained phenylalanine upregulation (Figs. 5, 6), suggests reinforcement of structural and antioxidant capacity. In maize, phenylalanine, tyrosine, and tryptophan biosynthesis pathways were significantly enriched under drought (Li et al., 2024), supporting the concept that aromatic amino acids function as hubs linking primary metabolism to stress-defense compounds. These coordinated adjustments indicate that recurrent drought reshaped central carbon–nitrogen balance.

Although our study did not assess transcript levels, the metabolic persistence observed across cycles and maintained in vegetatively derived propagules strongly suggests coordinated regulatory control. In maize, combined transcriptomic–metabolomic analysis revealed that drought-responsive metabolites are tightly associated with hub genes controlling amino acid and hormone-related pathways (Li et al., 2024). The parallel enrichment of amino acid pathways in both species reinforces the interpretation that recurrent drought induces systemic reprogramming of central metabolism rather than isolated metabolite accumulation. Propagules derived from drought-stressed plants maintained a more coordinated metabolic response under renewed drought compared with propagules derived from well-watered plants. The persistence of this metabolic responsiveness across vegetative propagation strongly supports the existence of drought-induced metabolic imprinting in sugarcane, indicating that prior stress exposure reshapes subsequent physiological and metabolic adjustment.

### 4.3 Root allocation reveals whole-plant integration of stress memory

Recurrent drought induced coordinated structural adjustments at the whole-plant level. During Phase I, root dry mass increased significantly in both genotypes across successive drought cycles at both the tillering and maturation stages (Fig. 7e,g), indicating enhanced belowground investment. It is well-established that plants increase root growth in response to water deficit to improve water absorption (Yang et al., 2023). This strategy was likely employed by sugarcane plants subjected to drought cycles as a mechanism to cope with water deficit. Rice plants exposed to recurrent drought events under nutrient-deficient conditions also exhibited increased root investment, as indicated by a 69% increase in the root/shoot dry mass ratio (Galviz et al., 2025). Marcos et al. (2018a) similarly reported greater root dry mass in sugarcane subjected to repeated drought, accompanied by increased root H_2_O_2_ levels without evidence of oxidative damage, suggesting a signaling role for ROS in coordinating adaptive root development. Comparable patterns have been observed in other crops. In wheat, root biomass initially declined under drought; however, plants with prior drought exposure exhibited significantly greater root biomass during subsequent stress compared with non-conditioned plants (Wang et al., 2021). This indicates that stress history can enhance belowground resilience, supporting improved water uptake capacity under recurring drought. In contrast, shoot dry mass was generally maintained across cycles (Fig. 7a,c), suggesting that recurrent drought did not impose major penalties on aboveground biomass accumulation. Leaf area responses were genotype- and stage-dependent. Notably, plants of the high productivity genotype IACSP95-5000 exposed to drought during tillering exhibited a significant increase in leaf area compared with well-watered plants (Fig. 7i), whereas other genotype-stage combinations showed maintenance rather than enhancement of canopy expansion. The greater leaf area observed in IACSP95-5000 coincided with the attenuated reductions in *A* during D2 and D3 at tillering (Fig. 2a), suggesting that improved physiological performance under recurrent drought supported sustained or even enhanced canopy development in this genotype. This genotype-specific response is consistent with previous evidence of differential physiological plasticity in sugarcane. Marchiori et al. (2017) demonstrated that IACSP95-5000 exhibits greater physiological plasticity, particularly in gas exchange and photochemical traits, compared with other genotypes relying primarily on morphological adjustments. The present findings extend this concept by showing that recurrent drought not only elicited plastic responses but consolidated them into a memory-associated structural pattern in a genotype with intrinsically high biomass potential.

The simultaneous increase in root dry mass and maintenance or, in the case of IACSP95-5000 at tillering, enhancement of leaf area indicates preferential carbon allocation toward belowground structures without compromising canopy development. Such coordinated allocation suggests that improved recovery kinetics and stabilization of photosynthetic performance under recurrent drought enabled sustained carbon supply to both shoots and roots. Importantly, these structural adjustments were not restricted to directly stressed plants. In Phase II, propagules from both genotypes derived from drought-stressed plants and maintained under irrigation (T/W and M/W) accumulated greater shoot and root biomass compared with propagules derived from plants without prior drought exposure (W/W) (Fig. 7c,f), indicating persistence of altered growth trajectories beyond immediate stress conditions. Under renewed drought, propagules from drought-stressed plants exhibited less severe reductions in shoot dry mass, root dry mass, and leaf area compared with W/D, consistent with their reduced integrated physiological impairment (Fig. 2) and faster recovery kinetics (Figs. S3, S4).

Evidence from field conditions further strengthens this interpretation. Sugarcane propagules derived from drought-exposed parental plants displayed increased stalk yield, shoot biomass, leaf area index and a larger root system compared with propagules from non-stressed plants (Pissolato et al., 2025). The convergence between controlled-environment and field evidence indicates that drought memory in sugarcane is not merely a transient physiological adjustment but a persistent whole-plant reorganization affecting biomass allocation, canopy architecture and yield components. Epigenetic mechanisms have been implicated in stress-induced modifications of root morphology in clonal and sexually reproduced species (Bian et al., 2013; Cortijo et al., 2014; Nosalewicz et al., 2016), suggesting a potential regulatory basis for persistent structural reprogramming. Enhanced root growth may improve water and nutrient uptake efficiency, supporting higher stomatal conductance and increased CO_2_ assimilation – a pattern consistent with the physiological data observed in the present study.

### 4.4 A proposed framework: drought-induced metabolic imprinting in clonal sugarcane

Taken together, our findings support a conceptual framework in which the first drought cycle acts as a critical reprogramming event rather than merely a transient stress episode. During D1, strong reductions in RWC, *A*_n_, *g*_s_, k_c_ and photochemical efficiency marked a threshold-level disturbance of carbon-water balance. This initial perturbation appears to initiate coordinated physiological and metabolic reorganization that is not fully reversed upon rehydration. Instead of resetting to a “pre-stress state”, plants retained an altered functional configuration. In subsequent drought cycles (D2 and D3), this preconditioned network was reactivated more efficiently, resulting in attenuated reductions in gas exchange, improved WUE_i_, preservation of photochemical performance, reinforced root allocation, and accelerated recovery kinetics. Importantly, this response was not restricted to immediate physiological regulation; it was accompanied by sustained reorganization of primary metabolism, including enhanced amino acid accumulation, modulation of TCA intermediates, and activation of the shikimate pathway.

In clonally derived propagules, this reprogrammed state was partially maintained. Propagules originating from drought-stressed plants exhibited lower integrated reductions in *A*_n_ and *g*_s_, faster restoration of carbon assimilation and carboxylation efficiency during rehydration, and enhanced biomass performance relative to propagules derived from non-conditioned plants. The persistence of altered responsiveness across vegetative propagation strongly suggests that the memory phenotype extends beyond transient metabolic adjustments. Because sugarcane propagation occurs through vegetative buds containing meristematic tissues (Dodd and Douhovnikoff, 2016), this persistence of drought responsiveness in clonally derived propagules suggests that stress-related regulatory information may be retained within bud meristems during vegetative propagation. Such transmission does not imply genetic inheritance but rather the maintenance of physiological or epigenetic states acquired during previous stress exposure (Pecinka et al., 2020; Gallusci et al., 2023; Latzel et al., 2016). In perennial and clonally propagated species, meristematic tissues may function as reservoirs of stress information, allowing conditioned physiological states to be transmitted to newly established plants derived from these buds.

Although molecular mechanisms were not directly assessed, the stability of this phenotype across stress cycles and clonal propagation is consistent with regulatory memory processes described in other species. Stress-induced chromatin remodeling, shifts in DNA methylation, transcriptional priming of metabolic genes, and persistence of signaling proteins or metabolites in inactive pools have all been implicated in drought memory (Hilker et al., 2015; Crisp et al., 2016; Virlouvet et al., 2018). The recovery phase likely serves as a consolidation window during which stress-related information is integrated into long-term regulatory states.

Based on our integrative physiological, metabolic and growth data, we propose that recurrent drought in sugarcane establishes a metabolically imprinted state characterized by: (1) recalibrated coordination between stomatal regulation and carbon assimilation; (2) stabilized photochemical performance under repeated stress; (3) sustained amino acid-based osmotic buffering; (4) reconfiguration of central carbon-nitrogen metabolism; (5) reinforced root allocation and whole-plant carbon partitioning; (6) enhanced post-stress recovery capacity. Collectively, these coordinated adjustments define a drought memory phenotype in clonal sugarcane. Rather than simply increasing tolerance at maximum stress, this phenotype reshapes the trajectory of stress response and recovery, integrating physiological control, metabolic restructuring and developmental plasticity into a stable and persistent functional state maintained across vegetative propagation.

## 5. Conclusion

Our results demonstrate that recurrent drought reshapes the physiological and metabolic responses of sugarcane rather than simply triggering repeated stress suppression. The first drought cycle acted as a conditioning event that reprogrammed plant responses, leading to attenuated physiological disruption, improved intrinsic water use efficiency, stabilization of photochemical performance, and coordinated metabolic adjustments in subsequent drought cycles. Metabolomic analyses further revealed sustained reorganization of primary metabolism, including modulation of amino acids and central carbon pathways, supporting the establishment of a metabolically imprinted state under recurrent water deficit. These responses varied according to genotype and developmental stage, with the high-biomass genotype IACSP95-5000 exhibiting greater structural plasticity and the drought-tolerant genotype IACCTC07-8008 showing more stable physiological performance. Importantly, this drought-induced imprint was maintained in clonally derived propagules, which exhibited reduced cumulative physiological disruption and faster recovery of photosynthetic performance. Elucidating the molecular mechanisms underlying this stress memory will be an important next step and may provide new opportunities to enhance resilience and productivity in sugarcane cultivation under increasingly variable climatic conditions.

