## Supplementary Material for "SUGARCANE’S DROUGHT MEMORY LEGACY: HOW PAST STRESS SHAPES FUTURE RESILIENCE"

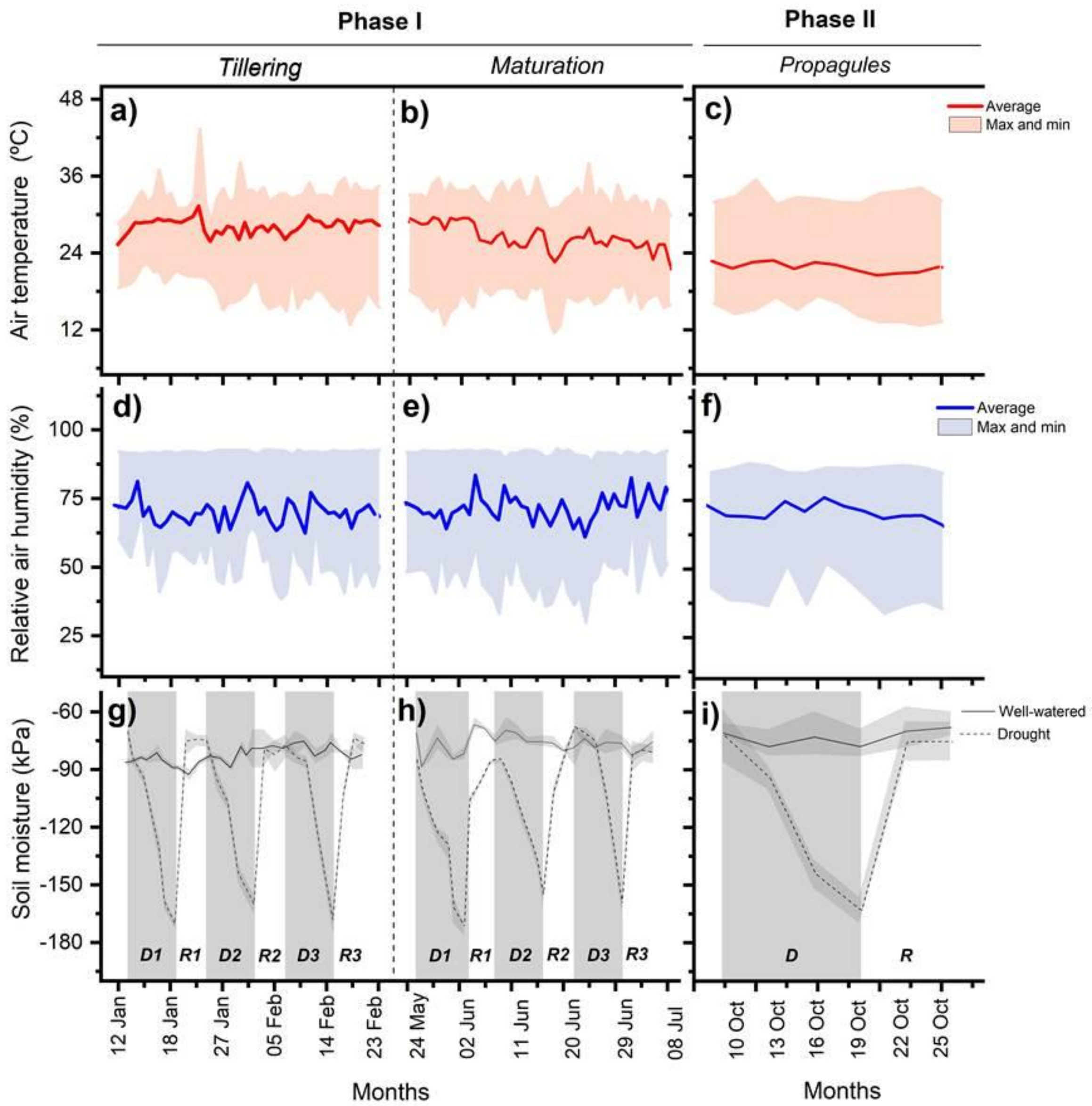

**Figure Supplementary 1.** Air temperature (average, maximum and minimum, in a-c), relative air humidity (average, maximum and minimum, in d-f) and soil moisture (in g-i) during the experimental period of phase I (origin material) and phase II (propagules). Shaded areas indicate drought cycles (D), and white areas indicate recovery periods (R). In soil moisture, solid and dashed lines represent well-watered and drought treatments, respectively. Each point represents the mean (n=6) ± SD.

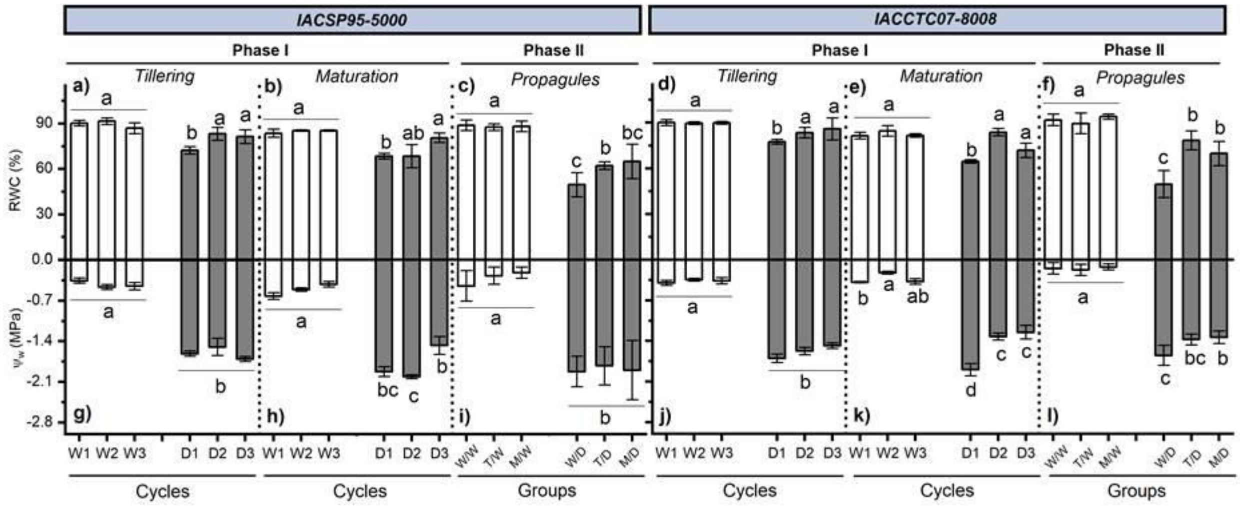

**Figure Supplementary 2.** Leaf water content (RWC, in a-f) and leaf water potential ($\Psi_W$, in g-l) of two sugarcane genotypes (IACSP95-5000 and IACCTC07-8008) subjected drought cycles (D) in phase I and phase II. In phase I plants were subjected to three drought cycles at the tillering or maturation stage, while phase II, the propagules derived from well-watered plants (W/) or from plants previously exposed to drought at tillering (T/) or maturation (M/) and subsequently subjected to either drought (/D) or kept well-hydrated (/W). The data were collected during the maximum water deficit (9$^{th}$ day). Bars represent mean values (n=4) ± SD. Different letters indicate significant differences among cycles or groups ($BF_{10}$ > 3).

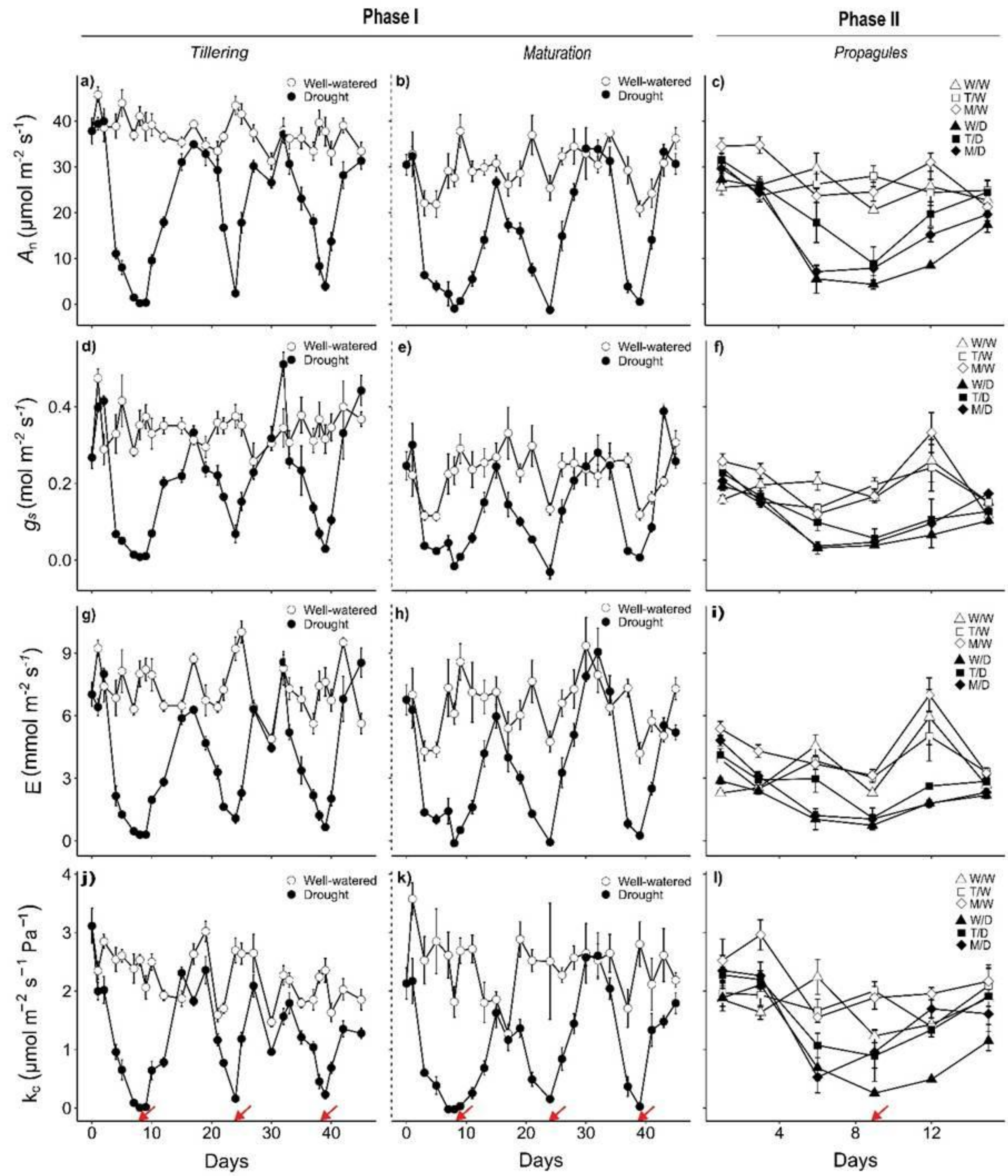

**Figure Supplementary 3**. Net $CO_2$ assimilation ($A_n$, in a-c), stomatal conductance ($g_s$, in d-f), transpiration (E, in g-i), and instantaneous carboxylation efficiency ($k_c$, in j-l) in the sugarcane genotype IACSP95-5000. Phase I depicts plants subjected to three drought cycles followed by recovery at the tillering or maturation stage. Phase II shows propagules derived from well-watered plants (W) or from plants previously exposed to drought cycles at the tillering (T/) or maturation (M/) stage, which were subsequently either subjected to drought (/D) or maintained under well-watered conditions (/W). Red arrows indicate the day of maximum water deficit ($9^{th}$ day). Symbols represent means (n = 4) ± SD.

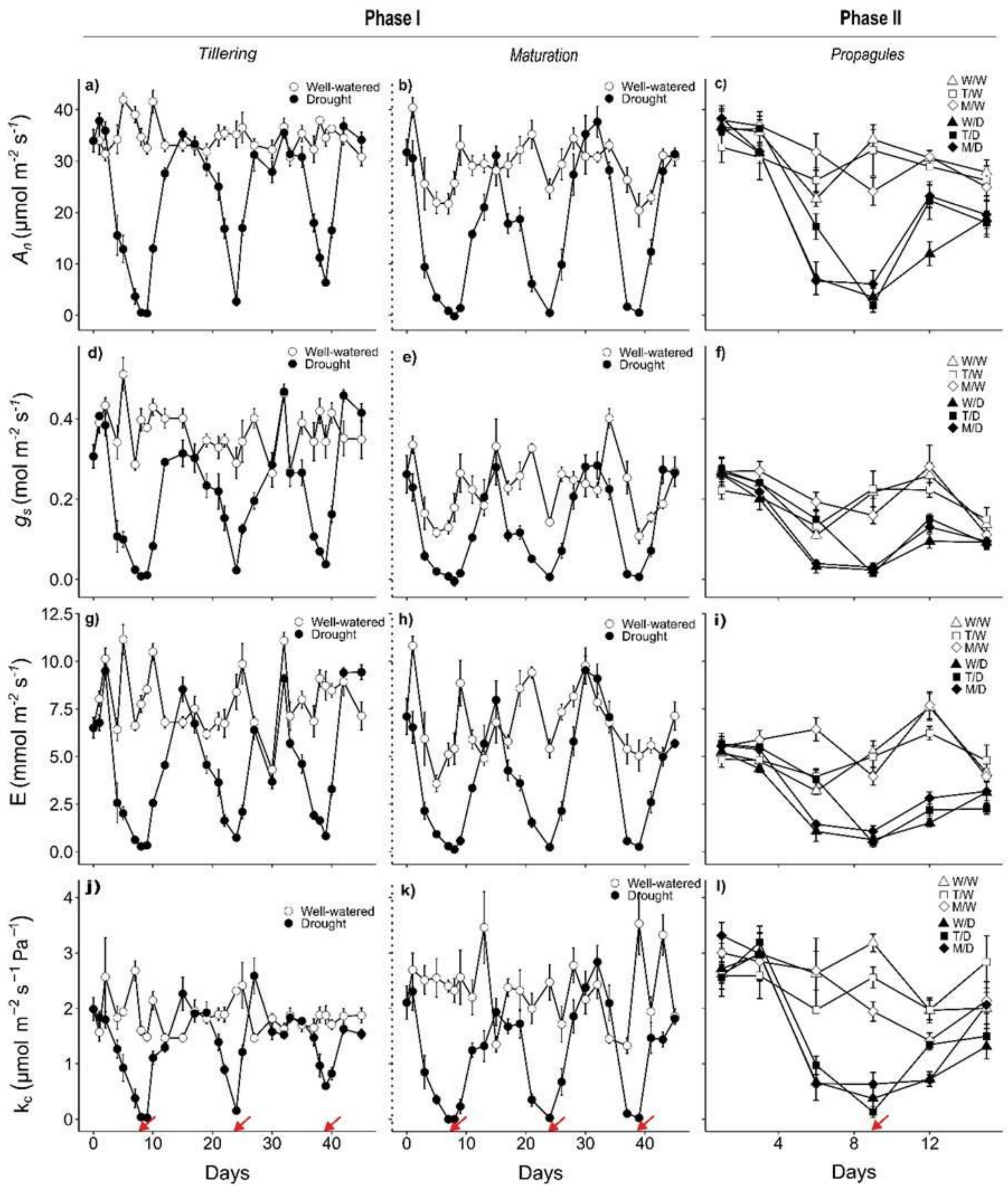

**Figure Supplementary 4**. Net $CO_2$ assimilation ($A_n$, in a-c), stomatal conductance ($g_s$, in d-f), transpiration (E, in g-i), and instantaneous carboxylation efficiency ($k_c$, in j-l) in the sugarcane genotype IACCTC07-8008. Phase I depicts plants subjected to three drought cycles (D) at the tillering (T) or maturation stage (M) or kept well-watered (W). Phase II shows sugarcane propagules derived from well-watered plants (W/) or from plants previously exposed to drought cycles at the tillering (T/) or maturation stage (M/), which were subsequently either subjected to drought (/D) or maintained under well-watered conditions (/W). Red arrows indicate the day of maximum water deficit (9$^{th}$ day). Symbols represent means (n=4) ± SD.

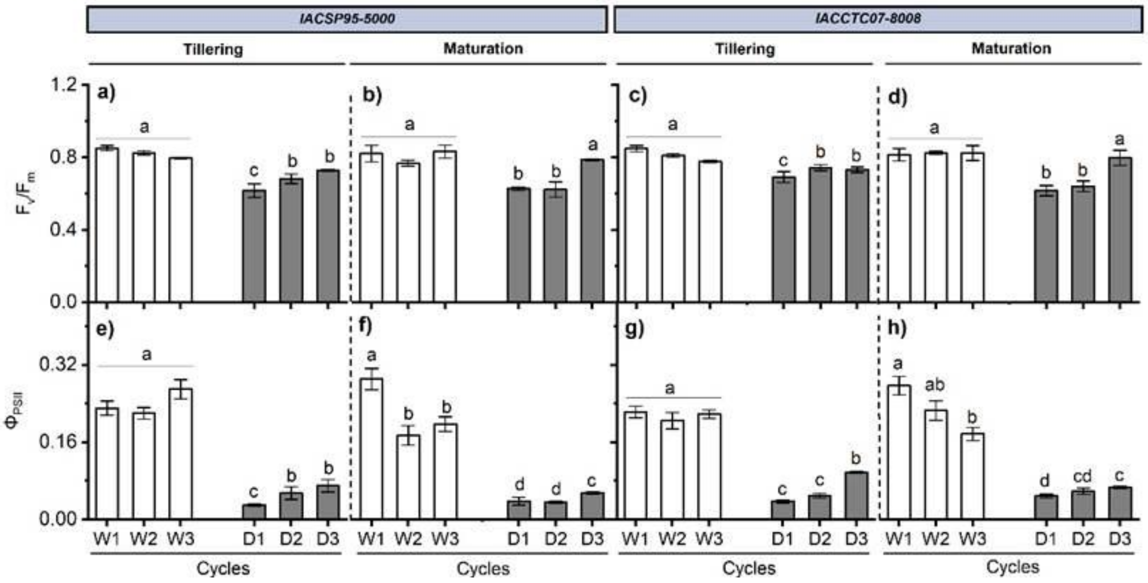

**Figure Supplementary 5.** Maximum quantum efficiency of PSII ($F_v/F_m$, in a-d) and effective quantum yield of PSII ($\Phi_{PSII}$, in e-h) of two sugarcane genotypes (IACSP95-5000 and IACCTC07-8008) subjected to three drought cycles (D) during tillering or maturation stage or kept well-watered (W). The data were collected during the maximum water deficit (9$^{th}$ day). Bars represent means (n=4) ± SD. Different letters indicate significant differences among cycles ($BF_{10}$> 3).

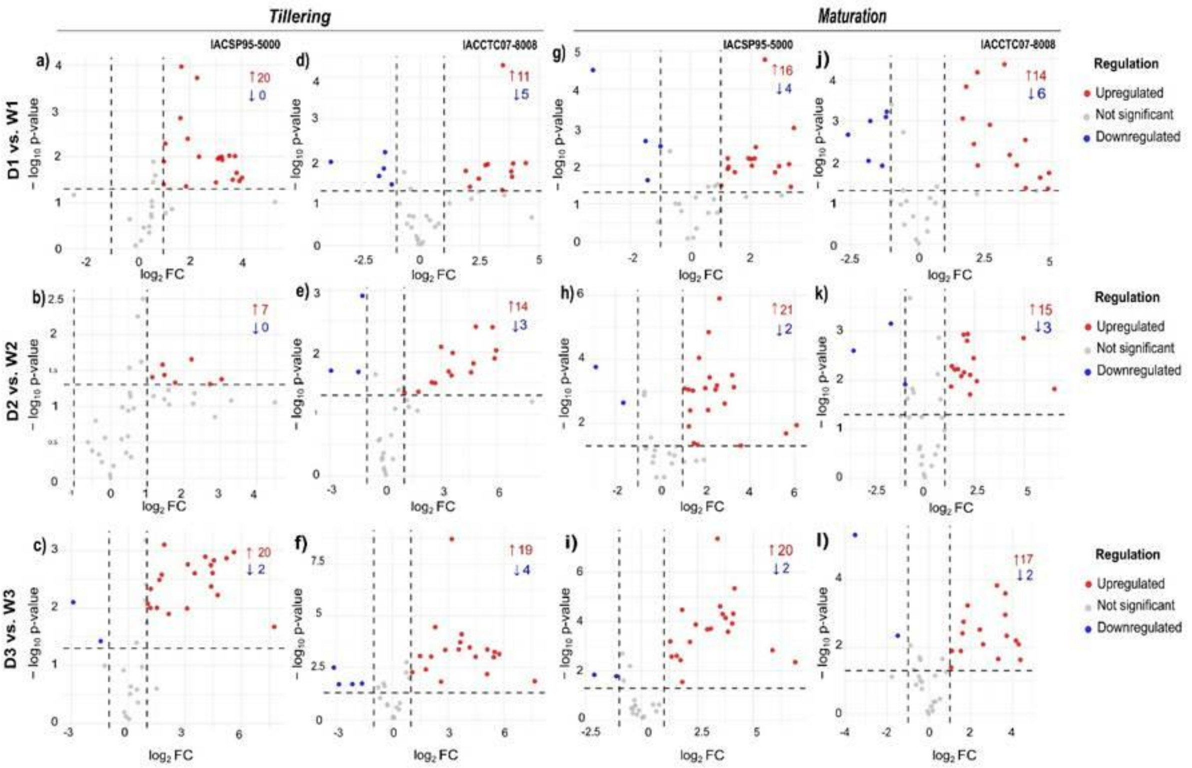

**Figure Supplementary 6.** Metabolite trends in two sugarcane genotypes (IACSP95-5000 and IACCTC07-8008) subjected to three drought cycles (D) during tillering or maturation or maintained under well-watered conditions (W). Volcano plots show differentially accumulated leaf primary metabolites in Phase I, comparing drought cycles (D1, D2 and D3) with their respective well-watered controls (W1, W2 and W3) at the tillering (a-f) and maturation (g-l) stages. Differential accumulation was assessed using two-sided Welch's t-tests. Metabolites with $p < 0.05$ and $\log_2 FC > 1$ (fold change > 2) are shown in red, whereas metabolites with $p < 0.05$ and $\log_2 FC < -1$ (fold change < 0.5) are shown in blue. Grey points represent metabolites that did not meet both significance criteria. Numbers indicate the total number of significantly increased and decreased metabolites.

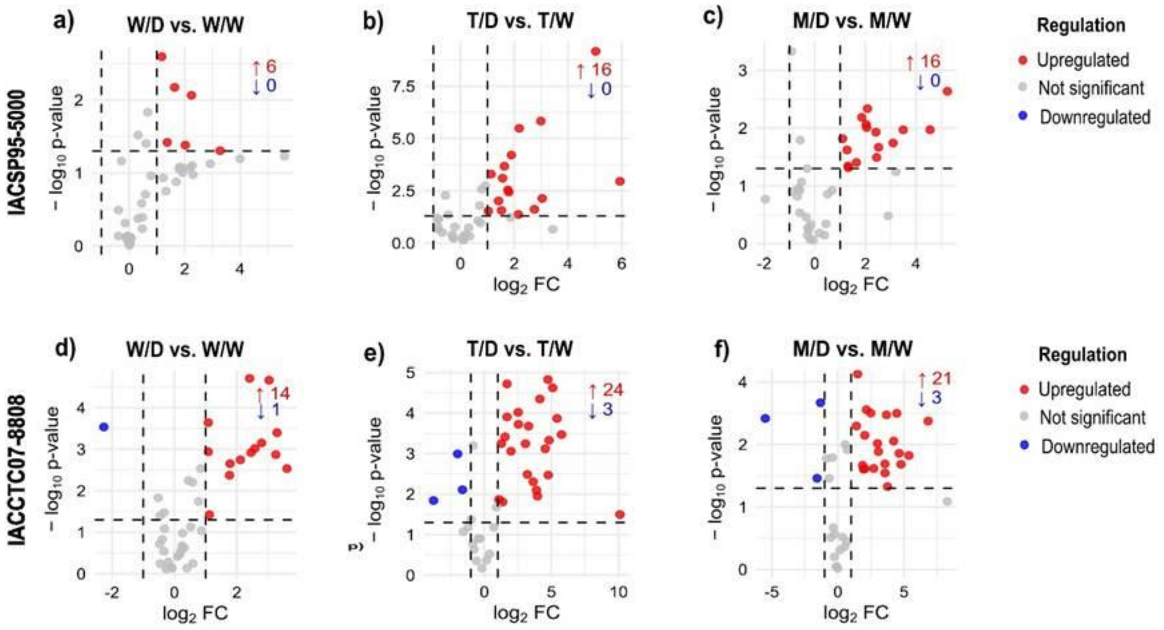

**Figure Supplementary 7.** Metabolite trends in two sugarcane genotypes (IACSP95-5000 and IACCTC07-8008) subjected to drought. Volcano plots show differentially accumulated leaf primary metabolites in Phase II. Propagules were obtained from well-watered plants (W/) or from plants previously exposed to drought during tillering (T/) or maturation (M/) and were subsequently either subjected to drought (/D) or maintained under well-watered conditions (/W). Differential accumulation was assessed using two-sided Welch's t-tests. Metabolites with $p < 0.05$ and $\log_2 FC > 1$ (fold change > 2) are shown in red, whereas metabolites with $p < 0.05$ and $\log_2 FC < -1$ (fold change < 0.5) are shown in blue. Grey points represent metabolites that did not meet both criteria. Numbers indicate the total number of significantly increased and decreased metabolites.

**Table supplementary 1.** Differentially accumulated metabolites in leaves of sugarcane genotype IACSP95-5000 subjected to drought cycles compared with well-watered plants during the tillering or maturation stages. For each metabolite, $\log_2$ fold change ($\log_2$FC) was calculated as the $\log_2$ ratio between mean metabolite abundance under drought and the corresponding well-watered control. Statistical significance was assessed using two-sided Welch's t-tests (n = 4). Only metabolites meeting both $p < 0.05$ and $|\log_2 FC| > 1$ were included. Positive $\log_2$FC values indicate increased abundance under drought, whereas negative values indicate decreased abundance.

| | PHENOLOGICAL PHASE | COMPARISON | METABOLITE | REGULATION | $Log_2FC$ | P-VALUE | PHENOLOGICAL PHASE | COMPARISON | METABOLITE | REGULATION | $Log_2FC$ | P-VALUE |
|---|---|---|---|---|---|---|---|---|---|---|---|---|
| **IACSP95-5000** | Tillering | D1 vs. W1 | β-Alanine | Upregulated | 3.02 | 0.036181 | Maturation | D1 vs. W1 | Aspartic acid | Downregulated | -1.43 | 0.024361 |
| **IACSP95-5000** | Tillering | D1 vs. W1 | Butanoate | Upregulated | 1.01 | 0.039661 | Maturation | D1 vs. W1 | Butanoate | Upregulated | 1.26 | 0.010924 |
| **IACSP95-5000** | Tillering | D1 vs. W1 | Fructose | Upregulated | 1.93 | 0.004058 | Maturation | D1 vs. W1 | Fructose | Upregulated | 2.47 | 1.70E-05 |
| **IACSP95-5000** | Tillering | D1 vs. W1 | Glucose (1MEOX) MP | Upregulated | 2.36 | 0.00998 | Maturation | D1 vs. W1 | Glucose (1MEOX) MP | Upregulated | 3.43 | 0.001049 |
| **IACSP95-5000** | Tillering | D1 vs. W1 | Glutamate | Upregulated | 2.29 | 1.93E-04 | Maturation | D1 vs. W1 | Glutarate | Downregulated | -3.24 | 3.20E-05 |
| **IACSP95-5000** | Tillering | D1 vs. W1 | Glycerol | Upregulated | 1.07 | 0.005173 | Maturation | D1 vs. W1 | Glycine | Upregulated | 1.23 | 0.012306 |
| **IACSP95-5000** | Tillering | D1 vs. W1 | Glycine | Upregulated | 3.91 | 0.03357 | Maturation | D1 vs. W1 | Isoleucine | Upregulated | 2.94 | 0.010479 |
| **IACSP95-5000** | Tillering | D1 vs. W1 | Isoleucine | Upregulated | 3.53 | 0.009455 | Maturation | D1 vs. W1 | Leucine | Upregulated | 2.02 | 0.00682 |
| **IACSP95-5000** | Tillering | D1 vs. W1 | Leucine | Upregulated | 3.15 | 0.010372 | Maturation | D1 vs. W1 | Lysine | Upregulated | 2.17 | 0.003319 |
| **IACSP95-5000** | Tillering | D1 vs. W1 | Lysine | Upregulated | 1.87 | 0.044341 | Maturation | D1 vs. W1 | Malate | Downregulated | -1.49 | 0.002307 |
| **IACSP95-5000** | Tillering | D1 vs. W1 | Methionine | Upregulated | 1.69 | 1.09E-04 | Maturation | D1 vs. W1 | Methionine | Upregulated | 1.23 | 0.006515 |
| **IACSP95-5000** | Tillering | D1 vs. W1 | Ornithine | Upregulated | 3.09 | 0.011088 | Maturation | D1 vs. W1 | Phenylalanine | Upregulated | 3.33 | 0.036284 |
| **IACSP95-5000** | Tillering | D1 vs. W1 | Phenylalanine | Upregulated | 3.81 | 0.02238 | Maturation | D1 vs. W1 | Proline | Upregulated | 3.27 | 0.009218 |
| **IACSP95-5000** | Tillering | D1 vs. W1 | Proline | Upregulated | 3.66 | 0.031668 | Maturation | D1 vs. W1 | Pyroglutamate | Upregulated | 2.04 | 0.01016 |
| **IACSP95-5000** | Tillering | D1 vs. W1 | Pyroglutamate | Upregulated | 3.75 | 0.009696 | Maturation | D1 vs. W1 | Quinic acid | Upregulated | 1.92 | 0.006525 |
| **IACSP95-5000** | Tillering | D1 vs. W1 | Serine | Upregulated | 4.01 | 0.028862 | Maturation | D1 vs. W1 | Serine | Upregulated | 2.81 | 0.015144 |
| **IACSP95-5000** | Tillering | D1 vs. W1 | Sugar unknown | Upregulated | 1.65 | 0.001434 | Maturation | D1 vs. W1 | Sugar unknown | Upregulated | 1.02 | 0.033507 |
| **IACSP95-5000** | Tillering | D1 vs. W1 | Threonate | Upregulated | 1.01 | 0.012695 | Maturation | D1 vs. W1 | Threonate | Downregulated | -1.00 | 0.003149 |
| **IACSP95-5000** | Tillering | D1 vs. W1 | Threonine | Upregulated | 3.24 | 0.010109 | Maturation | D1 vs. W1 | Threonine | Upregulated | 1.47 | 0.014967 |
| **IACSP95-5000** | Tillering | D1 vs. W1 | Valine | Upregulated | 3.25 | 0.011717 | Maturation | D1 vs. W1 | Valine | Upregulated | 2.13 | 0.006561 |
| **IACSP95-5000** | Tillering | D2 vs. W2 | β-Alanine | Upregulated | 1.14 | 0.039409 | Maturation | D2 vs. W2 | Adenine | Upregulated | 3.28 | 7.33E-04 |

| | | | | | | | | | | | | |
|---|---|---|---|---|---|---|---|---|---|---|---|---|
| **IACSP95-5000** | Tillering | D2 vs. W2 | Glutamate | Upregulated | 1.77 | 0.046814 | Maturation | D2 vs. W2 | β-Alanine | Upregulated | 3.21 | 3.10E-04 |
| **IACSP95-5000** | Tillering | D2 vs. W2 | Leucine | Upregulated | 3.04 | 0.042024 | Maturation | D2 vs. W2 | Asparagine | Upregulated | 6.09 | 0.011228 |
| **IACSP95-5000** | Tillering | D2 vs. W2 | Methionine | Upregulated | 1.47 | 0.036569 | Maturation | D2 vs. W2 | Butanoate | Upregulated | 1.33 | 0.003895 |
| **IACSP95-5000** | Tillering | D2 vs. W2 | Ornithine | Upregulated | 1.43 | 0.026294 | Maturation | D2 vs. W2 | Glutamate | Upregulated | 1.26 | 8.76E-04 |
| **IACSP95-5000** | Tillering | D2 vs. W2 | Threonine | Upregulated | 2.22 | 0.022198 | Maturation | D2 vs. W2 | Glutarate | Downregulated | -2.88 | 1.72E-04 |
| **IACSP95-5000** | Tillering | D2 vs. W2 | Valine | Upregulated | 2.72 | 0.049126 | Maturation | D2 vs. W2 | Glycerol | Upregulated | 1.28 | 0.012229 |
| **IACSP95-5000** | Tillering | D3 vs. W3 | Adenine | Upregulated | 1.92 | 7.94E-04 | Maturation | D2 vs. W2 | Glycine | Upregulated | 1.08 | 8.04E-04 |
| **IACSP95-5000** | Tillering | D3 vs. W3 | β-Alanine | Upregulated | 3.53 | 0.002487 | Maturation | D2 vs. W2 | Isoleucine | Upregulated | 1.73 | 9.01E-05 |
| **IACSP95-5000** | Tillering | D3 vs. W3 | Asparagine | Upregulated | 7.73 | 0.021193 | Maturation | D2 vs. W2 | Leucine | Upregulated | 1.46 | 9.39E-04 |
| **IACSP95-5000** | Tillering | D3 vs. W3 | 3-cis-caffeoylquinic acid | Downregulated | -1.45 | 0.037417 | Maturation | D2 vs. W2 | Lysine | Upregulated | 2.20 | 3.62E-04 |
| **IACSP95-5000** | Tillering | D3 vs. W3 | Glucose (1MEOX) MP | Upregulated | 1.66 | 0.003237 | Maturation | D2 vs. W2 | Malate | Downregulated | -1.65 | 0.002243 |
| **IACSP95-5000** | Tillering | D3 vs. W3 | Glutamate | Upregulated | 3.17 | 0.001745 | Maturation | D2 vs. W2 | Methionine | Upregulated | 2.46 | 8.78E-04 |
| **IACSP95-5000** | Tillering | D3 vs. W3 | Glutarate | Downregulated | -2.90 | 0.007937 | Maturation | D2 vs. W2 | Ornithine | Upregulated | 2.53 | 6.49E-04 |
| **IACSP95-5000** | Tillering | D3 vs. W3 | Glycerol | Upregulated | 1.18 | 0.009967 | Maturation | D2 vs. W2 | Phenylalanine | Upregulated | 2.64 | 1.27E-06 |
| **IACSP95-5000** | Tillering | D3 vs. W3 | Glycine | Upregulated | 3.13 | 0.010146 | Maturation | D2 vs. W2 | Proline | Upregulated | 5.61 | 0.020202 |
| **IACSP95-5000** | Tillering | D3 vs. W3 | Isoleucine | Upregulated | 5.21 | 0.001363 | Maturation | D2 vs. W2 | Pyroglutamate | Upregulated | 2.13 | 0.003823 |
| **IACSP95-5000** | Tillering | D3 vs. W3 | Leucine | Upregulated | 4.38 | 0.001801 | Maturation | D2 vs. W2 | Quinic acid | Upregulated | 3.60 | 0.048569 |
| **IACSP95-5000** | Tillering | D3 vs. W3 | Lysine | Upregulated | 1.53 | 0.009908 | Maturation | D2 vs. W2 | Serine | Upregulated | 2.88 | 0.002377 |
| **IACSP95-5000** | Tillering | D3 vs. W3 | Methionine | Upregulated | 1.05 | 0.008433 | Maturation | D2 vs. W2 | Sugar unknown | Upregulated | 1.66 | 0.045479 |
| **IACSP95-5000** | Tillering | D3 vs. W3 | Ornithine | Upregulated | 1.78 | 0.002629 | Maturation | D2 vs. W2 | Threonine | Upregulated | 2.15 | 1.42E-05 |
| **IACSP95-5000** | Tillering | D3 vs. W3 | Phenylalanine | Upregulated | 5.61 | 0.001068 | Maturation | D2 vs. W2 | 5-trans-caffeoylquinic acid | Upregulated | 1.49 | 0.040858 |
| **IACSP95-5000** | Tillering | D3 vs. W3 | Proline | Upregulated | 4.74 | 0.005958 | Maturation | D2 vs. W2 | Valine | Upregulated | 2.01 | 7.90E-04 |
| **IACSP95-5000** | Tillering | D3 vs. W3 | Pyroglutamate | Upregulated | 4.55 | 0.001488 | Maturation | D3 vs. W3 | Adenine | Upregulated | 1.57 | 0.002451 |
| **IACSP95-5000** | Tillering | D3 vs. W3 | Serine | Upregulated | 4.40 | 0.004222 | Maturation | D3 vs. W3 | β-Alanine | Upregulated | 3.87 | 2.62E-04 |
| **IACSP95-5000** | Tillering | D3 vs. W3 | Sugar unknown | Upregulated | 2.15 | 0.012633 | Maturation | D3 vs. W3 | Asparagine | Upregulated | 6.88 | 0.004389 |
| **IACSP95-5000** | Tillering | D3 vs. W3 | Threonate | Upregulated | 1.20 | 0.004627 | Maturation | D3 vs. W3 | Aspartic acid | Upregulated | 1.34 | 0.00264 |
| **IACSP95-5000** | Tillering | D3 vs. W3 | Threonine | Upregulated | 4.09 | 0.001286 | Maturation | D3 vs. W3 | Butanoate | Upregulated | 1.81 | 3.37E-05 |
| **IACSP95-5000** | Tillering | D3 vs. W3 | Valine | Upregulated | 4.48 | 0.002469 | Maturation | D3 vs. W3 | Glutamate | Upregulated | 2.43 | 1.32E-04 |
| | | | | | | | Maturation | D3 vs. W3 | Glutarate | Downregulated | -2.11 | 0.01438 |

| | | | | | | | |
|---|---|---|---|---|---|---|---|
| | | Maturation | D3 vs. W3 | Glycerol | Upregulated | 1.28 | 6.44E-04 |
| | | Maturation | D3 vs. W3 | Glycine | Upregulated | 2.17 | 6.67E-04 |
| | | Maturation | D3 vs. W3 | *Myo*-inositol | Upregulated | 1.74 | 0.003655 |
| | | Maturation | D3 vs. W3 | Isoleucine | Upregulated | 4.11 | 4.88E-05 |
| | | Maturation | D3 vs. W3 | Leucine | Upregulated | 4.16 | 4.59E-06 |
| | | Maturation | D3 vs. W3 | Lysine | Upregulated | 3.63 | 4.97E-05 |
| | | Maturation | D3 vs. W3 | Methionine | Upregulated | 3.10 | 1.95E-04 |
| | | Maturation | D3 vs. W3 | Ornithine | Upregulated | 1.82 | 0.027882 |
| | | Maturation | D3 vs. W3 | Phenylalanine | Upregulated | 3.40 | 4.46E-08 |
| | | Maturation | D3 vs. W3 | Proline | Upregulated | 5.86 | 0.001455 |
| | | Maturation | D3 vs. W3 | Pyroglutamate | Upregulated | 4.05 | 1.21E-04 |
| | | Maturation | D3 vs. W3 | Serine | Upregulated | 3.52 | 2.44E-05 |
| | | Maturation | D3 vs. W3 | Threonine | Upregulated | 2.94 | 2.09E-04 |
| | | Maturation | D3 vs. W3 | 4-trans-caffeoylquinic acid | Downregulated | -1.13 | 0.016406 |
| | | Maturation | D3 vs. W3 | Valine | Upregulated | 3.76 | 7.19E-05 |

**Table supplementary 2**. Differentially accumulated metabolites in leaves of sugarcane genotype IACCTC07-8008 subjected to drought cycles compared with well-watered plants during the tillering or maturation stages. For each metabolite, $\log_2$ fold change ($\log_2$FC) was calculated as the $\log_2$ ratio between mean metabolite abundance under drought and the corresponding well-watered control. Statistical significance was assessed using two-sided Welch's t-tests (n = 4). Only metabolites meeting both $p < 0.05$ and $|\log_2FC| > 1$ were included. Positive $\log_2$FC values indicate increased abundance under drought, whereas negative values indicate decreased abundance.

| CULTIVAR | PHENOLOGICAL PHASE | COMPARISON | METABOLITE | REGULATION | $Log_2FC$ | P-VALUE | PHENOLOGICAL PHASE | COMPARISON | METABOLITE | REGULATION | $Log_2FC$ | P-VALUE |
|---|---|---|---|---|---|---|---|---|---|---|---|---|
| **IACCTC07-8008** | Tillering | D1 vs. W1 | 3-cis-caffeoylquinic acid | Downregulated | -1.54 | 0.014628 | Maturation | D1 vs. W1 | Aspartic acid | Downregulated | -1.15 | 6.08E-04 |
| **IACCTC07-8008** | Tillering | D1 vs. W1 | Fructose | Upregulated | 3.47 | 5.14E-05 | Maturation | D1 vs. W1 | 3-cis-caffeoylquinic acid | Downregulated | -1.32 | 0.012428 |
| **IACCTC07-8008** | Tillering | D1 vs. W1 | Fumarate | Downregulated | -1.49 | 0.005958 | Maturation | D1 vs. W1 | Fructose | Upregulated | 2.23 | 6.70E-05 |
| **IACCTC07-8008** | Tillering | D1 vs. W1 | Glucose (1MEOX) MP | Upregulated | 2.83 | 0.011536 | Maturation | D1 vs. W1 | Fumarate | Downregulated | -1.18 | 8.12E-04 |
| **IACCTC07-8008** | Tillering | D1 vs. W1 | Glutamate | Upregulated | 1.91 | 0.016762 | Maturation | D1 vs. W1 | Glucose (1MEOX) MP | Upregulated | 3.24 | 4.30E-05 |
| **IACCTC07-8008** | Tillering | D1 vs. W1 | Glutarate | Upregulated | -3.77 | 0.010204 | Maturation | D1 vs. W1 | Glutamate | Upregulated | 1.68 | 8.85E-04 |
| **IACCTC07-8008** | Tillering | D1 vs. W1 | Glycine | Upregulated | 3.91 | 0.011149 | Maturation | D1 vs. W1 | Glutarate | Downregulated | -2.59 | 0.002189 |
| **IACCTC07-8008** | Tillering | D1 vs. W1 | Isoleucine | Upregulated | 3.86 | 0.023036 | Maturation | D1 vs. W1 | Glycine | Upregulated | 2.69 | 0.00126 |
| **IACCTC07-8008** | Tillering | D1 vs. W1 | Leucine | Upregulated | 3.83 | 0.017173 | Maturation | D1 vs. W1 | Isoleucine | Upregulated | 4.89 | 0.01857 |
| **IACCTC07-8008** | Tillering | D1 vs. W1 | Lysine | Upregulated | 2.09 | 0.039943 | Maturation | D1 vs. W1 | Leucine | Upregulated | 4.02 | 0.043548 |
| **IACCTC07-8008** | Tillering | D1 vs. W1 | Phenylalanine | Upregulated | 3.45 | 0.047576 | Maturation | D1 vs. W1 | Malate | Downregulated | -1.77 | 0.001021 |
| **IACCTC07-8008** | Tillering | D1 vs. W1 | Phosphoric acid | Downregulated | -1.21 | 0.034902 | Maturation | D1 vs. W1 | Methionine | Upregulated | 2.09 | 0.00369 |
| **IACCTC07-8008** | Tillering | D1 vs. W1 | Serine | Upregulated | 4.44 | 0.010857 | Maturation | D1 vs. W1 | Ornithine | Upregulated | 1.81 | 1.49E-04 |
| **IACCTC07-8008** | Tillering | D1 vs. W1 | Threonine | Upregulated | 2.46 | 0.02564 | Maturation | D1 vs. W1 | Phenylalanine | Upregulated | 4.56 | 0.024273 |
| **IACCTC07-8008** | Tillering | D1 vs. W1 | 3-trans-caffeoylquinic acid | Downregulated | -1.74 | 0.022121 | Maturation | D1 vs. W1 | Proline | Upregulated | 4.86 | 0.044726 |
| **IACCTC07-8008** | Tillering | D1 vs. W1 | Valine | Upregulated | 2.74 | 0.012146 | Maturation | D1 vs. W1 | Pyroglutamate | Upregulated | 3.43 | 0.006774 |
| **IACCTC07-8008** | Tillering | D2 vs. W2 | β-Alanine | Upregulated | 2.49 | 0.030907 | Maturation | D1 vs. W1 | Serine | Upregulated | 4.01 | 0.002937 |
| **IACCTC07-8008** | Tillering | D2 vs. W2 | 3-cis-caffeoylquinic acid | Downregulated | -1.25 | 0.001241 | Maturation | D1 vs. W1 | Threonine | Upregulated | 2.24 | 0.012188 |
| **IACCTC07-8008** | Tillering | D2 vs. W2 | Glutamate | Upregulated | 2.99 | 0.008241 | Maturation | D1 vs. W1 | 3-trans-caffeoylquinic acid | Downregulated | -1.83 | 0.00944 |
| **IACCTC07-8008** | Tillering | D2 vs. W2 | Glutarate | Downregulated | -2.94 | 0.020026 | Maturation | D1 vs. W1 | Valine | Upregulated | 3.70 | 0.012051 |
| **IACCTC07-8008** | Tillering | D2 vs. W2 | Glycine | Upregulated | 3.55 | 0.023877 | Maturation | D2 vs. W2 | Adenine | Upregulated | 2.26 | 0.019423 |

| | | | | | | | | | | | | |
|---|---|---|---|---|---|---|---|---|---|---|---|---|
| **IACCTC07-8008** | Tillering | D2 vs. W2 | *Myo*-inositol | Upregulated | 1.02 | 0.045588 | Maturation | D2 vs. W2 | β-Alanine | Upregulated | 2.44 | 0.003509 |
| **IACCTC07-8008** | Tillering | D2 vs. W2 | Isoleucine | Upregulated | 5.88 | 0.012674 | Maturation | D2 vs. W2 | Asparagine | Upregulated | 4.95 | 0.001357 |
| **IACCTC07-8008** | Tillering | D2 vs. W2 | Leucine | Upregulated | 5.96 | 0.0094 | Maturation | D2 vs. W2 | Aspartic acid | Downregulated | -1.72 | 6.91E-04 |
| **IACCTC07-8008** | Tillering | D2 vs. W2 | Lysine | Upregulated | 2.68 | 0.031745 | Maturation | D2 vs. W2 | 3-cis-caffeoylquinic acid | Downregulated | -1.00 | 0.012046 |
| **IACCTC07-8008** | Tillering | D2 vs. W2 | Methionine | Upregulated | 1.77 | 0.043996 | Maturation | D2 vs. W2 | Glutarate | Downregulated | -3.60 | 0.00247 |
| **IACCTC07-8008** | Tillering | D2 vs. W2 | Phenylalanine | Upregulated | 5.76 | 0.003986 | Maturation | D2 vs. W2 | Glycine | Upregulated | 2.27 | 0.007705 |
| **IACCTC07-8008** | Tillering | D2 vs. W2 | Proline | Upregulated | 4.86 | 0.003899 | Maturation | D2 vs. W2 | Isoleucine | Upregulated | 1.49 | 0.005972 |
| **IACCTC07-8008** | Tillering | D2 vs. W2 | Pyroglutamate | Upregulated | 3.62 | 0.01032 | Maturation | D2 vs. W2 | Leucine | Upregulated | 1.30 | 0.013371 |
| **IACCTC07-8008** | Tillering | D2 vs. W2 | Serine | Upregulated | 4.73 | 0.015231 | Maturation | D2 vs. W2 | Lysine | Upregulated | 1.95 | 0.006864 |
| **IACCTC07-8008** | Tillering | D2 vs. W2 | Threonine | Upregulated | 3.40 | 0.020598 | Maturation | D2 vs. W2 | Methionine | Upregulated | 2.15 | 0.001126 |
| **IACCTC07-8008** | Tillering | D2 vs. W2 | 3-trans-caffeoylquinic acid | Downregulated | -1.46 | 0.020893 | Maturation | D2 vs. W2 | Ornithine | Upregulated | 1.35 | 0.005113 |
| **IACCTC07-8008** | Tillering | D2 vs. W2 | Valine | Upregulated | 4.61 | 0.021221 | Maturation | D2 vs. W2 | Phenylalanine | Upregulated | 2.11 | 0.001551 |
| **IACCTC07-8008** | Tillering | D3 vs. W3 | β-Alanine | Upregulated | 3.58 | 4.13E-04 | Maturation | D2 vs. W2 | Proline | Upregulated | 6.50 | 0.014984 |
| **IACCTC07-8008** | Tillering | D3 vs. W3 | Asparagine | Upregulated | 7.67 | 0.013952 | Maturation | D2 vs. W2 | Pyroglutamate | Upregulated | 1.94 | 0.001155 |
| **IACCTC07-8008** | Tillering | D3 vs. W3 | Butanoate | Upregulated | 1.07 | 0.005189 | Maturation | D2 vs. W2 | Serine | Upregulated | 2.60 | 0.010357 |
| **IACCTC07-8008** | Tillering | D3 vs. W3 | 3-cis-caffeoylquinic acid | Downregulated | -2.15 | 0.019421 | Maturation | D2 vs. W2 | Threonine | Upregulated | 1.66 | 0.005896 |
| **IACCTC07-8008** | Tillering | D3 vs. W3 | Fructose | Upregulated | 2.09 | 9.18E-04 | Maturation | D2 vs. W2 | Valine | Upregulated | 1.82 | 0.008197 |
| **IACCTC07-8008** | Tillering | D3 vs. W3 | Glucose (1MEOX) MP | Upregulated | 2.62 | 0.014935 | Maturation | D3 vs. W3 | Adenine | Upregulated | 1.06 | 0.040123 |
| **IACCTC07-8008** | Tillering | D3 vs. W3 | Glutamate | Upregulated | 3.20 | 1.95E-09 | Maturation | D3 vs. W3 | β-Alanine | Upregulated | 2.59 | 0.008129 |
| **IACCTC07-8008** | Tillering | D3 vs. W3 | Glutarate | Downregulated | -3.15 | 0.003089 | Maturation | D3 vs. W3 | Asparagine | Upregulated | 4.29 | 0.008133 |
| **IACCTC07-8008** | Tillering | D3 vs. W3 | Glycine | Upregulated | 4.49 | 9.26E-04 | Maturation | D3 vs. W3 | Glucose (1MEOX) MP | Upregulated | 1.10 | 0.012669 |
| **IACCTC07-8008** | Tillering | D3 vs. W3 | Isoleucine | Upregulated | 5.76 | 6.87E-04 | Maturation | D3 vs. W3 | Glutamate | Upregulated | 1.53 | 0.012847 |
| **IACCTC07-8008** | Tillering | D3 vs. W3 | Leucine | Upregulated | 5.51 | 5.52E-04 | Maturation | D3 vs. W3 | Glutarate | Downregulated | -3.53 | 4.35E-06 |
| **IACCTC07-8008** | Tillering | D3 vs. W3 | Lysine | Upregulated | 2.85 | 4.32E-04 | Maturation | D3 vs. W3 | Glycine | Upregulated | 3.31 | 0.022384 |
| **IACCTC07-8008** | Tillering | D3 vs. W3 | Methionine | Upregulated | 1.81 | 0.003834 | Maturation | D3 vs. W3 | Isoleucine | Upregulated | 4.16 | 0.006333 |
| **IACCTC07-8008** | Tillering | D3 vs. W3 | Ornithine | Upregulated | 1.49 | 8.57E-04 | Maturation | D3 vs. W3 | Leucine | Upregulated | 4.37 | 0.023802 |
| **IACCTC07-8008** | Tillering | D3 vs. W3 | Phenylalanine | Upregulated | 5.46 | 9.75E-04 | Maturation | D3 vs. W3 | Lysine | Upregulated | 2.43 | 0.002953 |
| **IACCTC07-8008** | Tillering | D3 vs. W3 | Proline | Upregulated | 5.10 | 0.006371 | Maturation | D3 vs. W3 | Methionine | Upregulated | 1.61 | 0.003806 |
| **IACCTC07-8008** | Tillering | D3 vs. W3 | Pyroglutamate | Upregulated | 3.73 | 7.68E-05 | Maturation | D3 vs. W3 | Ornithine | Upregulated | 1.66 | 0.001721 |
| **IACCTC07-8008** | Tillering | D3 vs. W3 | Serine | Upregulated | 5.12 | 4.16E-04 | Maturation | D3 vs. W3 | Phenylalanine | Upregulated | 3.65 | 0.001065 |

| | | | | | | | | | | | | |
|---|---|---|---|---|---|---|---|---|---|---|---|---|
| **IACCTC07-8008** | Tillering | D3 vs. W3 | Sugar unknown | Upregulated | 2.31 | 3.44E-05 | Maturation | D3 vs. W3 | Proline | Upregulated | 6.08 | 0.035248 |
| **IACCTC07-8008** | Tillering | D3 vs. W3 | Threonine | Upregulated | 3.65 | 1.87E-04 | Maturation | D3 vs. W3 | Pyroglutamate | Upregulated | 3.25 | 1.41E-04 |
| **IACCTC07-8008** | Tillering | D3 vs. W3 | 3-trans-caffeoylquinic acid | Upregulated | -2.88 | 0.019747 | Maturation | D3 vs. W3 | Serine | Upregulated | 3.66 | 2.45E-04 |
| **IACCTC07-8008** | Tillering | D3 vs. W3 | 5-trans-caffeoylquinic acid | Upregulated | -1.63 | 0.018096 | Maturation | D3 vs. W3 | Threonine | Upregulated | 1.85 | 5.60E-04 |
| **IACCTC07-8008** | Tillering | D3 vs. W3 | Valine | Upregulated | 4.17 | 3.30E-04 | Maturation | D3 vs. W3 | 3-trans-caffeoylquinic acid | Downregulated | -1.48 | 0.004495 |
| | | | | | | | Maturation | D3 vs. W3 | Valine | Upregulated | 3.72 | 0.004525 |

**Table S3.** Differentially accumulated metabolites in leaves of sugarcane propagules of IACSP95-5000 and IACCTC07-8008 derived from well-watered plants or from plants previously exposed to drought during the tillering or maturation stages. Propagules were subsequently subjected to drought or maintained under well-watered conditions. For each metabolite, $log_2$ fold change ($log_2FC$) was calculated as the $log_2$ ratio between mean metabolite abundance under drought and the corresponding well-watered control. Statistical significance was assessed using two-sided Welch's t-tests (n = 4). Only metabolites meeting both $p < 0.05$ and $|log_2FC| > 1$ were included. Positive $log_2FC$ values indicate increased abundance under drought, whereas negative values indicate decreased abundance.

| CULTIVAR | COMPARISON | METABOLITE | REGULATION | $Log_2FC$ | P-VALUE | CULTIVAR | COMPARISON | METABOLITE | REGULATION | $Log_2FC$ | P-VALUE |
|---|---|---|---|---|---|---|---|---|---|---|---|
| **IACSP95-5000** | W/D vs W/W | Butanoate | Upregulated | 1.37 | 0.038 | **IACCTC07-8008** | W/D vs W/W | Asparagine | Upregulated | 3.03 | 2.17E-05 |
| **IACSP95-5000** | W/D vs W/W | Fructose | Upregulated | 1.17 | 0.003 | **IACCTC07-8008** | W/D vs W/W | β-Alanine | Upregulated | 1.08 | 2.28E-04 |
| **IACSP95-5000** | W/D vs W/W | Glucose (1MEOX) BP | Upregulated | 1.64 | 0.007 | **IACCTC07-8008** | W/D vs W/W | Fructose | Upregulated | 3.29 | 3.99E-04 |
| **IACSP95-5000** | W/D vs W/W | Glucose (1MEOX) MP | Upregulated | 2.25 | 0.009 | **IACCTC07-8008** | W/D vs W/W | Glucose (1MEOX) BP | Upregulated | 2.58 | 9.57E-04 |
| **IACSP95-5000** | W/D vs W/W | Glutarate | Upregulated | 2.02 | 0.042 | **IACCTC07-8008** | W/D vs W/W | Glucose (1MEOX) MP | Upregulated | 3.25 | 1.34E-03 |
| **IACSP95-5000** | W/D vs W/W | Ornithine | Upregulated | 3.27 | 0.049 | **IACCTC07-8008** | W/D vs W/W | Glutarate | Downregulated | -2.26 | 2.91E-04 |
| **IACSP95-5000** | T/D vs T/W | Asparagine | Upregulated | 5.93 | 0.001 | **IACCTC07-8008** | W/D vs W/W | Glycine | Upregulated | 2.11 | 1.80E-03 |
| **IACSP95-5000** | T/D vs T/W | β-Alanine | Upregulated | 1.53 | 0.028 | **IACCTC07-8008** | W/D vs W/W | Isoleucine | Upregulated | 2.45 | 1.20E-03 |
| **IACSP95-5000** | T/D vs T/W | Butanoate | Upregulated | 1.42 | 0.010 | **IACCTC07-8008** | W/D vs W/W | Ornithine | Upregulated | 1.12 | 3.79E-02 |
| **IACSP95-5000** | T/D vs T/W | Fructose | Upregulated | 2.15 | 0.043 | **IACCTC07-8008** | W/D vs W/W | Phenylalanine | Upregulated | 2.79 | 7.05E-04 |
| **IACSP95-5000** | T/D vs T/W | Glucose (1MEOX) MP | Upregulated | 2.75 | 0.024 | **IACCTC07-8008** | W/D vs W/W | Pyroglutamate | Upregulated | 1.08 | 1.15E-03 |
| **IACSP95-5000** | T/D vs T/W | Glycine | Upregulated | 1.04 | 0.029 | **IACCTC07-8008** | W/D vs W/W | Serine | Upregulated | 3.60 | 2.92E-03 |
| **IACSP95-5000** | T/D vs T/W | Isoleucine | Upregulated | 2.99 | 0.000 | **IACCTC07-8008** | W/D vs W/W | Threonine | Upregulated | 1.78 | 2.22E-03 |
| **IACSP95-5000** | T/D vs T/W | Leucine | Upregulated | 2.19 | 0.000 | **IACCTC07-8008** | W/D vs W/W | Valine | Upregulated | 1.76 | 4.22E-03 |
| **IACSP95-5000** | T/D vs T/W | Lysine | Upregulated | 1.89 | 0.000 | **IACCTC07-8008** | W/D vs W/W | Sugar unknown | Upregulated | 2.54 | 1.20E-05 |
| **IACSP95-5000** | T/D vs T/W | Methionine | Upregulated | 1.14 | 0.001 | **IACCTC07-8008** | T/D vs T/W | Adenine | Upregulated | 1.99 | 8.72E-04 |
| **IACSP95-5000** | T/D vs T/W | Ornithine | Upregulated | 3.04 | 0.007 | **IACCTC07-8008** | T/D vs T/W | Asparagine | Upregulated | 10.11 | 3.18E-02 |
| **IACSP95-5000** | T/D vs T/W | Phenylalanine | Upregulated | 5.03 | 0.000 | **IACCTC07-8008** | T/D vs T/W | β-Alanine | Upregulated | 3.96 | 1.11E-02 |
| **IACSP95-5000** | T/D vs T/W | Pyroglutamate | Upregulated | 1.80 | 0.004 | **IACCTC07-8008** | T/D vs T/W | Butanoate | Upregulated | 1.54 | 3.86E-04 |

| | | | | | | | | | | | |
|---|---|---|---|---|---|---|---|---|---|---|---|
| **IACSP95-5000** | T/D vs T/W | Serine | Upregulated | 1.76 | 0.003 | **IACCTC07-8008** | T/D vs T/W | Fructose | Upregulated | 1.69 | 1.25E-04 |
| **IACSP95-5000** | T/D vs T/W | Threonine | Upregulated | 1.57 | 0.001 | **IACCTC07-8008** | T/D vs T/W | Glucose (1MEOX) BP | Upregulated | 3.07 | 5.69E-04 |
| **IACSP95-5000** | T/D vs T/W | Valine | Upregulated | 1.63 | 0.000 | **IACCTC07-8008** | T/D vs T/W | Glucose (1MEOX) MP | Upregulated | 2.52 | 9.55E-05 |
| **IACSP95-5000** | M/D vs M/W | Asparagine | Upregulated | 5.23 | 0.002 | **IACCTC07-8008** | T/D vs T/W | Glutamate | Upregulated | 2.54 | 1.89E-04 |
| **IACSP95-5000** | M/D vs M/W | β-Alanine | Upregulated | 1.33 | 0.048 | **IACCTC07-8008** | T/D vs T/W | Glutarate | Downregulated | -3.78 | 1.44E-02 |
| **IACSP95-5000** | M/D vs M/W | Fructose | Upregulated | 1.31 | 0.045 | **IACCTC07-8008** | T/D vs T/W | Glycerate | Upregulated | 1.37 | 1.56E-02 |
| **IACSP95-5000** | M/D vs M/W | Glucose (1MEOX) BP | Upregulated | 1.64 | 0.039 | **IACCTC07-8008** | T/D vs T/W | Glycerol | Upregulated | 1.09 | 1.35E-02 |
| **IACSP95-5000** | M/D vs M/W | Glucose (1MEOX) MP | Upregulated | 2.52 | 0.022 | **IACCTC07-8008** | T/D vs T/W | Glycine | Upregulated | 3.30 | 2.10E-04 |
| **IACSP95-5000** | M/D vs M/W | Isoleucine | Upregulated | 3.49 | 0.011 | **IACCTC07-8008** | T/D vs T/W | Isoleucine | Upregulated | 4.74 | 1.49E-05 |
| **IACSP95-5000** | M/D vs M/W | Leucine | Upregulated | 2.44 | 0.032 | **IACCTC07-8008** | T/D vs T/W | Leucine | Upregulated | 3.88 | 7.96E-03 |
| **IACSP95-5000** | M/D vs M/W | Lysine | Upregulated | 1.85 | 0.007 | **IACCTC07-8008** | T/D vs T/W | Lysine | Upregulated | 3.64 | 4.95E-03 |
| **IACSP95-5000** | M/D vs M/W | Methionine | Upregulated | 1.28 | 0.024 | **IACCTC07-8008** | T/D vs T/W | Malate | Downregulated | -1.99 | 1.02E-03 |
| **IACSP95-5000** | M/D vs M/W | Ornithine | Upregulated | 2.41 | 0.012 | **IACCTC07-8008** | T/D vs T/W | Methionine | Upregulated | 3.21 | 3.25E-03 |
| **IACSP95-5000** | M/D vs M/W | Phenylalanine | Upregulated | 4.55 | 0.011 | **IACCTC07-8008** | T/D vs T/W | Ornithine | Upregulated | 4.76 | 3.38E-03 |
| **IACSP95-5000** | M/D vs M/W | Proline | Upregulated | 1.10 | 0.015 | **IACCTC07-8008** | T/D vs T/W | Phenylalanine | Upregulated | 5.42 | 1.35E-04 |
| **IACSP95-5000** | M/D vs M/W | Pyroglutamate | Upregulated | 2.03 | 0.009 | **IACCTC07-8008** | T/D vs T/W | Phosphoric acid | Upregulated | 1.27 | 5.65E-04 |
| **IACSP95-5000** | M/D vs M/W | Serine | Upregulated | 3.09 | 0.018 | **IACCTC07-8008** | T/D vs T/W | Proline | Upregulated | 5.75 | 3.37E-04 |
| **IACSP95-5000** | M/D vs M/W | Threonine | Upregulated | 2.08 | 0.005 | **IACCTC07-8008** | T/D vs T/W | Pyroglutamate | Upregulated | 4.82 | 4.69E-04 |
| **IACSP95-5000** | M/D vs M/W | Valine | Upregulated | 2.05 | 0.010 | **IACCTC07-8008** | T/D vs T/W | Serine | Upregulated | 5.10 | 2.39E-05 |
| | | | | | | **IACCTC07-8008** | T/D vs T/W | Sucrose | Downregulated | -1.63 | 7.77E-03 |
| | | | | | | **IACCTC07-8008** | T/D vs T/W | Threonine | Upregulated | 4.14 | 4.48E-05 |
| | | | | | | **IACCTC07-8008** | T/D vs T/W | Valine | Upregulated | 4.54 | 7.58E-04 |
| | | | | | | **IACCTC07-8008** | T/D vs T/W | Sugar unknown | Upregulated | 1.51 | 1.20E-05 |
| | | | | | | **IACCTC07-8008** | M/D vs M/W | Adenine | Upregulated | 2.15 | 2.77E-03 |
| | | | | | | **IACCTC07-8008** | M/D vs M/W | β-Alanine | Upregulated | 1.86 | 2.15E-02 |
| | | | | | | **IACCTC07-8008** | M/D vs M/W | Fructose | Upregulated | 2.99 | 9.62E-03 |
| | | | | | | **IACCTC07-8008** | M/D vs M/W | Glucose (1MEOX) BP | Upregulated | 3.67 | 3.36E-03 |
| | | | | | | **IACCTC07-8008** | M/D vs M/W | Glucose (1MEOX) MP | Upregulated | 2.49 | 3.14E-03 |
| | | | | | | **IACCTC07-8008** | M/D vs M/W | Glutamate | Upregulated | 2.05 | 2.45E-02 |
| | | | | | | **IACCTC07-8008** | M/D vs M/W | Glutarate | Downregulated | -5.49 | 3.82E-03 |

| | | | | | | | |
|---|---|---|---|---|---|---|---|
| | | **IACCTC07-8008** | M/D vs M/W | Glycerate | Upregulated | 2.03 | 7.12E-03 |
| | | **IACCTC07-8008** | M/D vs M/W | Glycine | Upregulated | 4.45 | 3.17E-03 |
| | | **IACCTC07-8008** | M/D vs M/W | Isoleucine | Upregulated | 4.77 | 2.08E-02 |
| | | **IACCTC07-8008** | M/D vs M/W | Leucine | Upregulated | 3.75 | 4.72E-02 |
| | | **IACCTC07-8008** | M/D vs M/W | Lysine | Upregulated | 3.06 | 1.30E-02 |
| | | **IACCTC07-8008** | M/D vs M/W | Malate | Downregulated | -1.11 | 1.05E-03 |
| | | **IACCTC07-8008** | M/D vs M/W | Methionine | Upregulated | 1.92 | 2.48E-02 |
| | | **IACCTC07-8008** | M/D vs M/W | *Myo*-inositol | Upregulated | 1.40 | 5.06E-03 |
| | | **IACCTC07-8008** | M/D vs M/W | Ornithine | Upregulated | 2.71 | 2.42E-02 |
| | | **IACCTC07-8008** | M/D vs M/W | Phenylalanine | Upregulated | 5.37 | 1.50E-02 |
| | | **IACCTC07-8008** | M/D vs M/W | Proline | Upregulated | 5.86 | 9.16E-04 |
| | | **IACCTC07-8008** | M/D vs M/W | Pyroglutamate | Upregulated | 3.56 | 2.86E-02 |
| | | **IACCTC07-8008** | M/D vs M/W | Serine | Upregulated | 4.22 | 8.82E-03 |
| | | **IACCTC07-8008** | M/D vs M/W | Threonine | Upregulated | 3.56 | 2.04E-02 |
| | | **IACCTC07-8008** | M/D vs M/W | Urea | Downregulated | -1.58 | 3.48E-02 |
| | | **IACCTC07-8008** | M/D vs M/W | Valine | Upregulated | 4.63 | 1.38E-02 |
| | | **IACCTC07-8008** | M/D vs M/W | Sugar unknown | Upregulated | 1.10 | 1.20E-05 |
